# Quartet DNA reference materials and datasets for comprehensively evaluating germline variants calling performance

**DOI:** 10.1101/2022.09.28.509844

**Authors:** Luyao Ren, Xiaoke Duan, Lianghua Dong, Rui Zhang, Jingcheng Yang, Yuechen Gao, Rongxue Peng, Wanwan Hou, Yaqing Liu, Jingjing Li, Ying Yu, Naixin Zhang, Jun Shang, Fan Liang, Depeng Wang, Hui Chen, Lele Sun, Lingtong Hao, The Quartet Project Team, Andreas Scherer, Jessica Nordlund, Wenming Xiao, Joshua Xu, Weida Tong, Xin Hu, Peng Jia, Kai Ye, Jinming Li, Li Jin, Leming Shi, Huixiao Hong, Jing Wang, Shaohua Fan, Xiang Fang, Yuanting Zheng

## Abstract

Current methods for evaluating the accuracy of germline variant calls are restricted to easy-to-detect high-confidence regions, thus ignoring a substantial portion of difficult variants beyond the benchmark regions. We established four DNA reference materials from immortalized cell lines derived from a Chinese Quartet including parents and monozygotic twins. We integrated benchmark calls of 4.2 million small variants and 15,000 structural variants from multiple platforms and bioinformatic pipelines for evaluating the reliability of germline variant calls inside the benchmark regions. The genetic built-in-truth of the Quartet family design not only improved sensitivity of benchmark calls by removing additional false positive variants with apparently high quality, but also enabled estimation of the precision of variants calls outside the benchmark regions. Batch effects of variant calling in large-scale DNA sequencing efforts can be effectively identified with the concurrent use of the Quartet DNA reference materials along with study samples, and can be alleviated by training a machine learning model with the Quartet reference datasets to remove potential artifact calls. Matched RNA and protein reference materials were also established in the Quartet project, thereby enabling benchmark calls constructed from DNA reference materials for evaluation of variants calling performance on RNA and protein data. The Quartet DNA reference materials from this study are a resource for objective and comprehensive assessment of the accuracy of germline variant calls throughout the whole-genome regions.

## Introduction

The detection of germline variants from high-throughput DNA sequencing (DNA-seq) is vital for biomedical research and molecular diagnostics of rare^1^ and complex^2^ genetic diseases. Well-characterized genomic reference materials can be used to benchmark measurement procedures, calibrate measuring systems and determine flagging criteria, and thereby support reliable application of genomic sequencing in basic research and clinical practice^3, 4^.

Genome in a Bottle (GIAB) and other efforts have established various whole-genome reference materials and defined benchmark calls and regions to benchmark germline small variants (SNVs and indels)^5–8^ and structural variants (SVs)^9–11^. However, all these efforts on genomic reference materials only evaluated variants identified inside the benchmark regions. The full extent of sequences generated and analyzed for a test genome is greater than what is defined by the boundaries of the benchmark regions. A substantial portion of variants detected outside the benchmark regions are overlooked, including many medically relevant variants^12^. Moreover, benchmark calls and regions are generally integrated from various sequencing technologies and bioinformatic pipelines, and thus biased toward easy-to-detect genomic contexts. Using variants calling performance inside the benchmark regions as a proxy will overestimate the overall performance of DNA assays or bioinformatic pipelines on the whole-genome region. Moreover, ignoring variants outside the benchmark regions will militate against objective understanding of the limitations of existing sequencing technologies, and thus hindering further method development.

Furthermore, in many practical applications of omics technologies, especially in large cohort studies, samples are often inevitably processed by multiple sequencing platforms at multiple centers over a relatively long period of time^13^. These large-scale projects usually suffer from batch effects due to the inconsistency of experimental conditions and sequencing machines^14, 15^. In DNA sequencing, batch effects are largely overlooked, but their widespread existence could lead to incorrectly taking batch-specific artifacts as real biological findings. Genomic reference materials are effective tools to identify and mitigate batch effects in DNA-seq^16^. Genomic reference materials can be sequenced along with test samples in every batch to determine whether batch effects exist. According to the properties of true positives and false positives detected from genomic reference materials, proper thresholds can be selected to remove batch-specific artifacts for each batch^17^.

To address these challenges in DNA-seq and beyond, we established four DNA reference materials from Epstein Barr Virus (EBV)-immortalized lymphoblastoid cell lines of a Chinese Quartet family, including the biological parents and monozygotic twin daughters. The Quartet was recruited from the Fudan Taizhou cohort in Central China, possessing genetic features of both Northern and Southern Chinese populations^18^. We extensively sequenced the whole genomes of the Quartet reference samples using multiple short-read and long-read sequencing platforms. We integrated both small variant and structural variant benchmark sets for each of the Quartet reference samples for evaluating variants calling accuracy inside the benchmark regions. The genomes of the monozygotic twins are almost identical^19^, and the expected number of germline *de novo* variants is fewer than 30 per generation and fewer than 1000 somatic mutations are introduced from cell culture^20^. The number of Mendelian violations in the detected variants is far more than the expected numbers of germline *de novo* variants and somatic mutations, indicating that most of the violations are sequencing or calling errors. Pedigree information of the Quartet members not only helped improve the sensitivity of benchmark sets by eliminating additional false positive variants with apparently high quality, but also facilitated the estimation of false positive rates of variants called outside the benchmark regions. The diverse sequencing data from the Quartet DNA reference materials also allowed us to identify batch effects present in whole-genome sequencing (WGS). The Quartet pedigree information was used to develop a machine learning based batch-specific filtration strategy to remove false positives and improve cross-batch reproducibility.

This study is part of the Quartet Project that aims for quality control and data integration of multiomic profiling (http://chinese-quartet.org/). Apart from the DNA reference materials, the Quartet Project also established matched RNA, protein and metabolite reference materials from the same culturing of the immortalized Quartet cell lines. Benchmark sets defined for the DNA reference materials facilitate evaluation of variants calling accuracy from RNA and protein data according to the principles of the central dogma. Accompanying papers on the overall project findings[Zheng], transcriptomics[Yu], proteomics[Tian], metabolomics[Zhang], batch-effect monitoring and correction[Yu], and the Quartet Data Portal[Yang] can be found elsewhere^21–26^.

## Results

### Study design with monozygotic twins and data generation

We established four immortalized lymphoblastoid cell lines of a Chinese Quartet family, including father (F7), mother (M8), and monozygotic twin daughters (D5 and D6) (Fig. 1a). The Quartet DNA reference materials are genomic DNA (gDNA) extracted from each immortalized lymphoblastoid cell line in large single batches. To unbiasedly characterize germline small variants and SV benchmark calls, we sequenced all four Quartet genomes on four short-read (Illumina HiSeq and NovaSeq, BGI MGISEQ-2000 and DNBSEQ-T7 (30-60x coverage)) and three long-read (Oxford Nanopore Technologies (ONT) (100x coverage), Pacific Biosciences (PacBio) Sequel (80x coverage), and PacBio Sequel II (30x coverage)) sequencing platforms at seven centers. We then used four orthogonal technologies, including linked-read sequencing (10x Genomics (30x coverage)), SNP array (the Axiom Precision Medicine Research Array (PMRA)), optical sequencing (BioNano), and PacBio circular consensus sequencing (CCS) reads (50x coverage) to validate and refine the benchmark calls (Fig. 1a and Supplementary Table 1).

**Figure 1.**
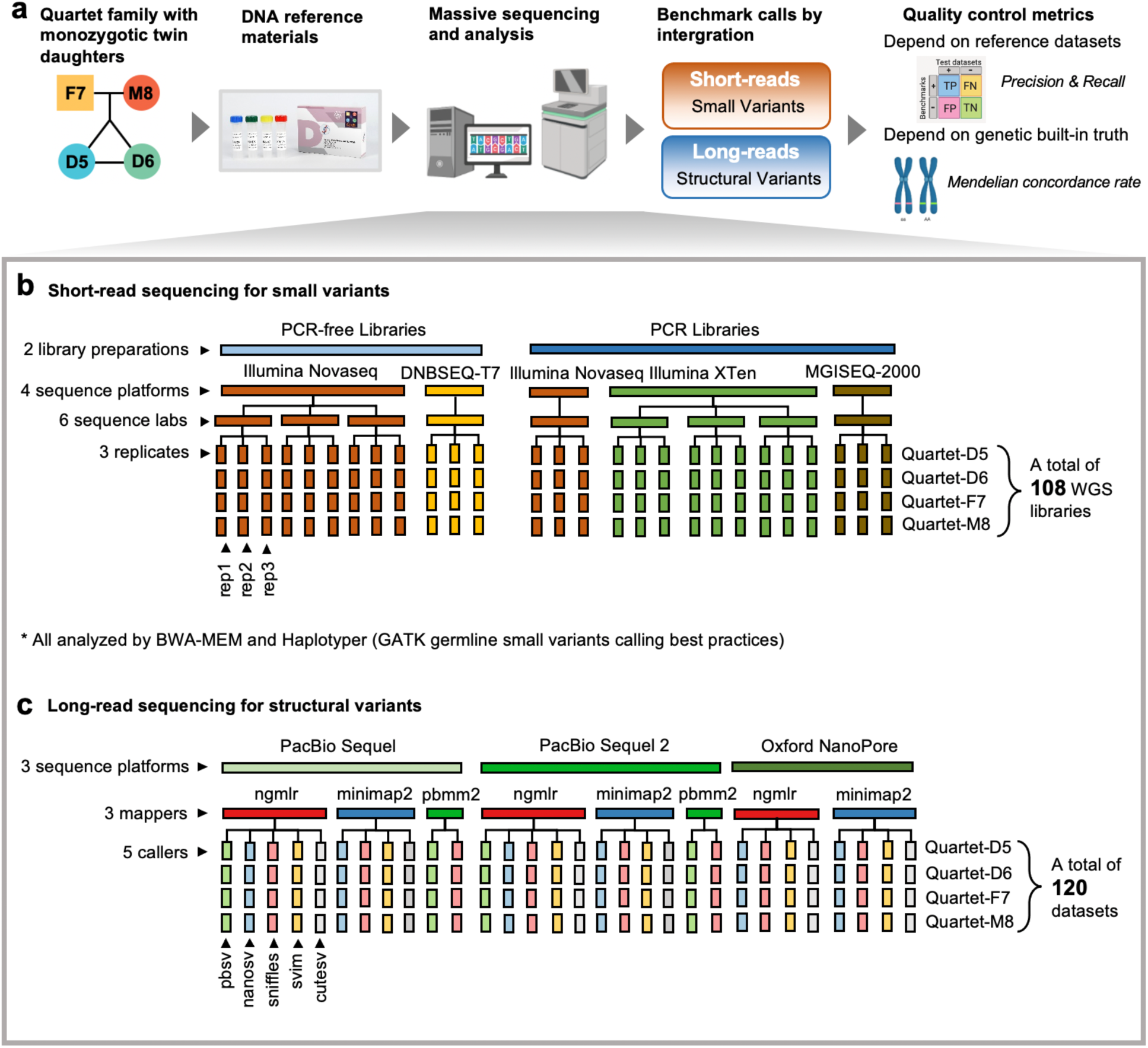
Study design and data generation. **(a)** Overview of the study design. Briefly, DNA reference materials were constructed from immortalized cell lines of a Chinese Quartet with father (F7), mother (M8), and monozygotic twin daughters (D5 and D6). They were sequenced by four short- and three long-read platforms at seven labs. Small variant and structural variant benchmark calls were integrated from massive sequencing datasets. Performance of a test dataset can be evaluated by comparing with benchmark calls or genetic built-in truth within the Quartet family. **(b)** Schematic overview of short-read sequencing datasets. Three replicates for each of the Quartet DNA reference materials were sequenced in nine batches, by both PCR and PCR-free libraries on four sequencing platforms at six labs, resulting in 108 WGS libraries. **(c)** Schematic overview of long-read sequencing datasets. One replicate for each of the Quartet DNA reference materials was sequenced per batch by PacBio Sequel, PacBio Sequel II and ONT. Eleven combinations of three mappers and five callers were used to detect structural variants, resulting in 120 variants calling datasets.

A total of 108 germline small variants call sets were obtained from 27 short-read WGS libraries of each Quartet genome using the widely adopted GATK best practices (BWA-MEM and HaplotypeCaller (HC)) (Fig. 1b and Supplementary Table 2). A total of 120 germline SV call sets were obtained from three long-read WGS libraries of each Quartet genome with 11 combinations from three aligners (NGMLR^27^, minimap2^28^, and pbmm2) and five callers (Sniffles^27^, NanoSV^29^, cuteSV^30^, SVIM^31^, and pbsv) (Fig. 1c and Supplementary Tables 3 and 4).

Variants call sets of the monozygotic twins are expected to be the same, because the twins share the identical genome from their parents. When investigating the consistency of call sets from different sequencing platforms, variants calling methods, and Quartet samples (Supplementary Fig. 1), we observed that SNVs, small indels (<50 bp), large insertions, or large deletions (>50 bp), were clustered distinctly based on the identity of the Quartet samples, and the monozygotic twins were grouped together as expected. However, for large duplications, inversions, or translocations (>50 bp), the call sets did not cluster by the identity of the Quartet samples, but revealed strong clustering by bioinformatic pipelines, indicating lack of reliability of or consistency in bioinformatic pipelines for these three types of SVs. Thus, these three types of SVs were not included in the benchmark sets.

### Determining small variant benchmark calls and regions

To define germline variant benchmark calls, we first selected reproducible variants among call sets for each of the Quartet samples. Because the number of Mendelian violations was much more than the expected number of *de novo* mutations or somatic mutations arising from cell culture, all Mendelian violations were assumed to be errors^20^. Thus, we excluded Mendelian violations from the benchmark calls, even when they were reproducible among call sets.

We generated one small variant benchmark dataset by integrating 108 call sets (27 call sets per sample) of all four Quartet samples based on short-read WGS. At the individual sample level, we obtained a total of 6 million variants of 27 call sets at the beginning, and an average of ∼4.6 million consensus variants after voting across triplicates in a batch, sequencing labs, and library preparation methods (PCR-free and PCR) (Fig. 2). To check Mendelian consistency of the remaining variants, genotypes should be confidently detected in all four Quartet samples for each variant. We then removed a total of 412,054 variant positions with no-call or conflict genotypes among the 27 call sets of any Quartet sample. Compared with variants filtered during the voting process, these removed variants showed higher variant allele frequency (VAF), read depth, mapping quality, and genotype quality (Supplementary Fig. 2). Therefore, they could not be removed by simply increasing variants filtration thresholds.

**Figure 2.**
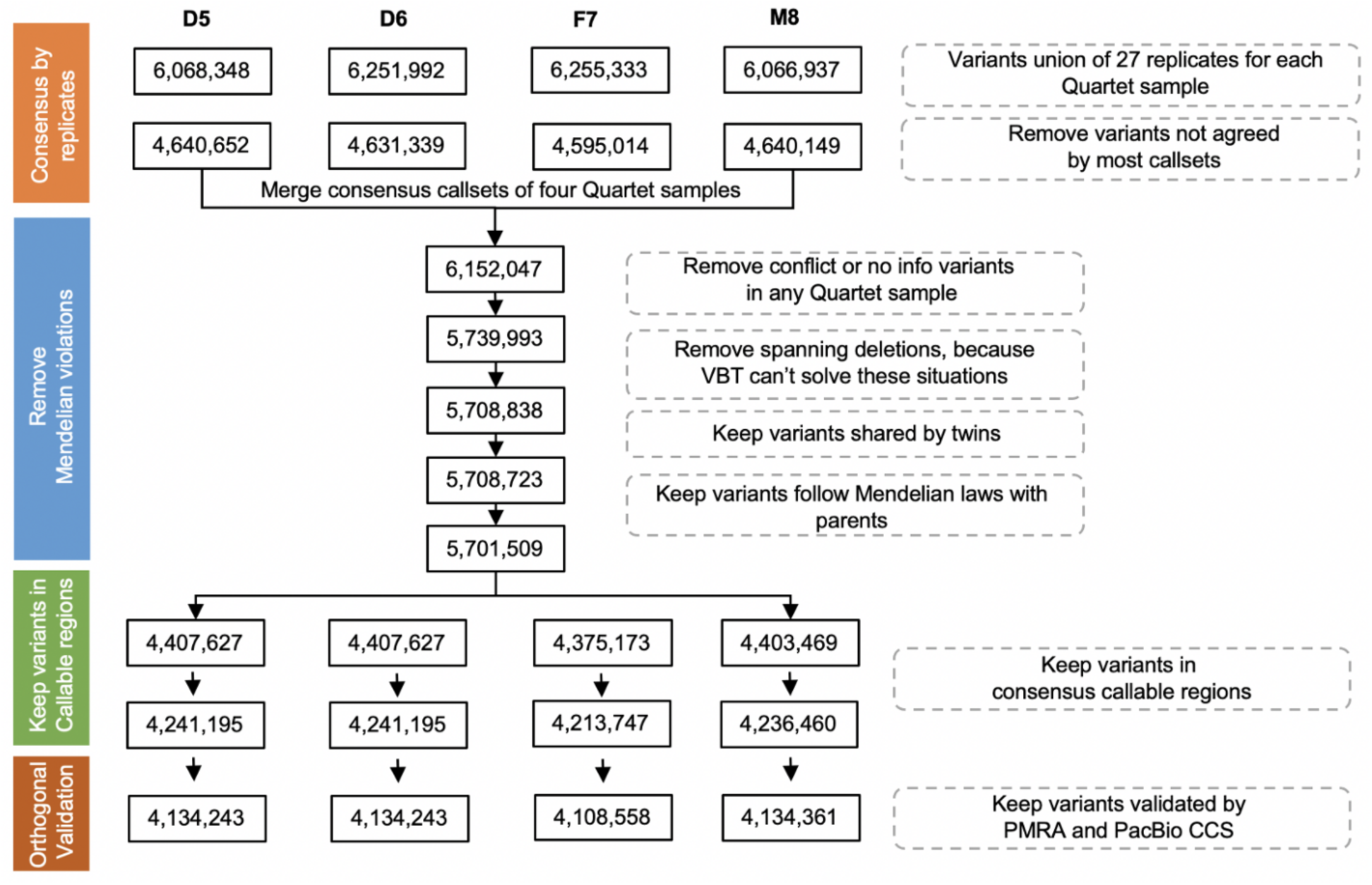
Integration workflow of Quartet small variant and structural variant benchmark calls. This workflow depicted the integration process to obtain small variant benchmark calls from 108 original GVCF call sets. Numbers in the boxes represented remaining small variants after each data processing step in the grey dotted boxes. Approximately 6 million small variants were discovered in 27 call sets for each Quartet reference sample. About 1.5 million small variants were removed by the voting process. We merged the four consensus call sets corresponding to the four Quartet samples, and discarded variants that did not reach agreement across 27 replicates in any Quartet sample. Only Mendelian consistent variants, which were shared by twins and following Mendelian inheritance laws and validated by PMRA and PacBio CCS datasets, were kept as small variant benchmark calls.

We identified 5,708,723 small variant positions with reproducible genotype calls among all four Quartet samples. These remaining variants were further examined for Mendelian consistency in the Quartet family, and 7,329 (0.13%) of them were identified as Mendelian violations. We manually inspected 4,761 variants located in the callable regions with high mapping quality. Of the 3,221 validated small variants, 1034 overlapped with large deletions. They were mistakenly considered as Mendelian discordant by Mendelian analysis software VBT^32^, which was based on the hypothesis that variants always passed on diploid. Comparing with the variants detected in the matched blood samples of the Quartet family members, we found 95 pre-twinning germline *de novo* variants shared by the twins (homozygous reference in the parents and heterozygous or homozygous alternative in the twins), one postzygotic germline *de novo* variant specifically found in Quartet-D5, 1,532 somatic variants (also found in blood), and 559 variants probably accumulated from cell culture (not found in blood) (Supplementary Table 5). Finally, we kept the Mendelian violations confirmed by manually curation into the initial catalog of benchmark calls. This process resulted in about 4.2 million well-supported small variants for each Quartet sample.

Previous studies show that PacBio CCS reads yield a higher variant calling accuracy compared with NGS, especially when calling variants in the repetitive regions of the genome. When comparing with the variants based on 50x coverage of PacBio CCS reads, we found that 98.7% and 95.0% of the variants in our benchmark dataset can be validated (Supplementary Table 6). The 89.7% unvalidated ones were found to be located in the repetitive regions of the genome, especially segmental duplications (41.6%) and centromeres regions (27.9%).

We also validated the small variant benchmark dataset using 16 replicates of PMRA SNP array. We obtained 793,024 Mendelian consistent probes that were well-supported by most replicates from the 902,394 clinically related probes assayed on the PMRA array. Of those reliable probes, 99.99% homozygotic references, 98.6% SNVs, 95.7% small insertions, and 96.2% small deletions were the same with the NGS consensus variants (Supplementary Table 7). Among the 2,845 discordant variants, 2,704 were detected by the PMRA array but were absent from NGS. We manually inspected the read alignment and found that the remaining 141 calls were either missed by NGS or genotyped differently from the PMRA array, and only seven were obvious false positive in the NGS consensus calls due to misalignment of NGS reads. The seven obvious false positives were later removed from the small variant benchmark calls. Consequently, the two validation processes removed 61,532 SNVs and 61,152 indels from the benchmark call sets.

To enable the identification of false positive and false negative variants, we defined benchmark regions for small variants (Supplementary Fig. 3). Benchmark regions were defined as high-confidence variant and homozygotic reference regions in consensus callable regions by all Quartet samples, covering 87.8% of GRCh38 reference genome (∼2.66 G; chr1-22, X). Consensus and Mendelian consistent variants outside the benchmark regions were not included in the final benchmark call sets (Table 1).

**Table 1.** Summary of Quartet small variant and structural variant benchmark calls and regions.

|  |  | Quartet-D5 | Quartet-D6 | Quartet-F7 | Quartet-M8 |
| --- | --- | --- | --- | --- | --- |
| Small Variant<br>Benchmark Calls | Total variants <sup>1</sup> | 4,134,243 | 4,134,243 | 4,108,558 | 4,134,361 |
|  | SNV | 3,564,768 | 3,564,768 | 3,544,173 | 3,564,170 |
|  | sINS <sup>2</sup> | 278,747 | 278,747 | 276,996 | 280,237 |
|  | sDEL <sup>2</sup> | 281,938 | 281,938 | 279,177 | 281,495 |
|  | Block Substitutions <sup>3</sup> | 8,790 | 8,790 | 8212 | 8,459 |
|  | Het/Hom Ratio | 1.37 | 1.37 | 1.30 | 1.35 |
|  | SNV Ti/Tv | 2.08 | 2.08 | 2.07 | 2.07 |
|  | Benchmark region,<br>chr1-22, X(bp) | 2,662,213,268 | 2,662,213,268 | 2,662,213,268 | 2,662,213,268 |
| Structural Variants<br>Benchmark Calls | Total Variants <sup>4</sup> | 15,005 | 15,005 | 15,098 | 14,893 |
|  | INS <sup>5</sup> < 1kb | 7,216 | 7,216 | 7,353 | 7,161 |
|  | INS ≥ 1kb | 734 | 734 | 755 | 717 |
|  | DEL <sup>5</sup> < 1kb | 6,352 | 6,352 | 6,287 | 6,324 |
|  | DEL ≥ 1kb | 703 | 703 | 703 | 691 |
|  | Het/Hom Ratio | 1.43 | 1.43 | 1.45 | 1.57 |
|  | Longest INS (bp) | 12,450 | 12,450 | 12,450 | 12,450 |
|  | Longest DEL (bp) | 117,310 | 117,310 | 435,343 | 494,712 |
|  | Affected bases (bp) | 7,796,176 | 7,796,176 | 8,821,558 | 8,550,650 |
|  | Benchmark region,<br>chr1-22 (bp) | 2,622,728,511 | 2,622,728,511 | 2,591,967,148 | 2,596,140,552 |
| <sup>1</sup> All small variants benchmark calls are located in small variants high-confidence region, and false positive variants detected by orthogonal validation have been removed;<br><sup>2</sup> sINS and sDEL stand for short insertion and deletion with size less than 50 bp;<br><sup>3</sup> Block Substitutions are variants with length change between REF and ALT. It is not simple addition or removal of bases, for example, ATT -> CTTT;<br><sup>4</sup> Structural variants benchmark calls include variants not located in benchmark regions;<br><sup>5</sup> INS and DEL stand for long insertion and deletion with size over 50 bp. |  |  |  |  |  |

We further compared the small variants benchmark calls with high confidence call sets from two accompanying studies^33, 34^ (Supplementary Fig. 4). These two high confidence call sets provide orthogonal confirmation of our calls (FDU), since Pan et al. (NCTR) integrated high confidence calls from four mappers (Bowtie2, BWA, ISAAC, and Stampy) and six callers (FreeBayes, GATK-HC, ISAAC, Samtools, SNVer, and Varscan), and Jia et al. (XJTU) constructed haplotype-resolved high confidence calls by combining short-read and long-read technologies. We compared variants in the intersect of the three high confidence regions of the three studies and found that 99.99% SNVs and 99.51% indels in our FDU callset could be confirmed by either the NCTR callset or the XJTU callset.

### Determining structural variant benchmark calls and regions

A similar strategy was used to determine SV benchmark calls by integrating the 120 call sets obtained from the long-read WGS data (Fig. 3). Because a large SV may be incorrectly called as multiple adjacent smaller SVs, we clustered SVs of the same type within 1 kb in each call set. This left about 90,000 isolated SVs of each Quartet sample. Then, SVs supported by the same pipeline from at least two sequencing platforms or by at least six pipelines from the same platform were determined as consensus SVs. Large SVs over 10 Mb and the ones located in centromeres, peri-centromere, and gaps regions of the reference genome were excluded. The remaining 31,659 SVs were then re-genotyped in a pedigree using three genotypers (Sniffles^27^, SVjedi^35^, and LRcaller^36^) with the reads of PacBio Sequel and ONT. Consensus genotypes (23,891) from at least six of the ten genotype call sets were then determined as the consensus genotype calls for each of the Quartet samples. SVs with conflict genotypes had higher VAF (0.12-0.25 and 0.75-1.0) compared with discordant variants among replicates (0.12), but not as high as VAFs at peaks near 0.5 (heterozygous) or 1.0 (homozygous), respectively (Supplementary Fig. 5).

**Figure 3.**
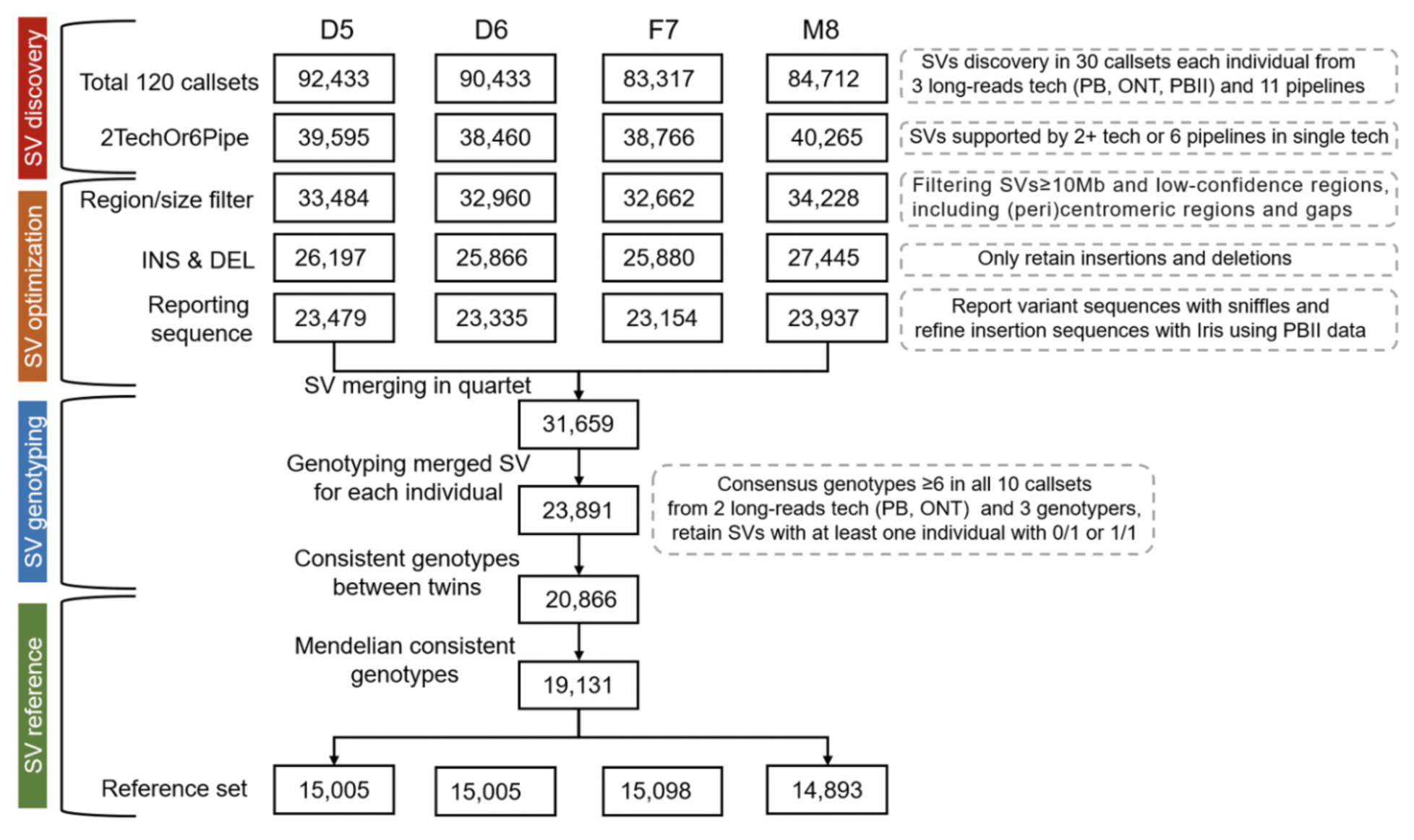
Integration workflow of structural variant benchmark calls. This workflow depicts the integration process to obtain structural variants benchmark calls from 120 call sets. Numbers in the box represented remaining structural variants after each data processing step in the grey dotted boxes. Briefly, approximately 90,000 structural variants were discovered in 30 call sets of each Quartet reference sample. We first kept structural variants supported by at least two sequencing platforms or at least six pipelines from one sequencing platforms, then removed SVs with length over 10 Mb or located on centromeric or peri-centromeric regions and gaps. INSs and DELs were extracted for the construction of structural variants benchmark calls. Sniffles was used to report structural variants sequences, and structural variants that failed in reporting sequences were filtered. Iris was applied to refine variant sequences. After obtaining consensus of structural variants in multiple data sets, we merged four catalogs of reproducible variants of each Quartet reference sample and obtained 31,659 SVs in total. Three genotypers were used to determine genotypes of these SVs, and only SVs with consensus genotypes in at least six of all ten genotype call sets were kept for Mendelian analysis. The final structural variants benchmark calls were shared by twins and followed Mendelian inheritance laws with parents.

After obtaining consensus genotyped SVs, we then removed Mendelian violated SVs. Of the 194 Mendelian violated SVs, we found that an 1,820 bp germline *de novo* heterozygous deletion shared by the twins, one specific heterozygous deletion of a twin daughter and two homozygous deletions of the father which probably were somatic or arose from cell culturing (Supplementary Table 8). These four SVs were supported by all three long-read platforms and thus retained in the benchmark call set. Following manual curation, we observed that the remaining 190 SVs were incorrectly genotyped. Most of them (91.7%) were located in regions of simple repeats over 100 bp or segmental duplications, or clustered with other variants. Therefore, these 190 SVs were not included in the benchmark call set. Finally, ∼15,000 benchmark SVs were kept into the benchmark call set for each Quartet sample (Table 1). Consistent with prior studies, we observed three peaks near 300 bp, 2.1 kb and 6 kb, likely reflecting the activities of *Alu* elements, SVA elements, and full-length LINE1 elements in the human genome (Supplementary Fig. 6).

Validating based on Illumina short reads, 10X Genomics linked-read, BioNano optical mapping, and whole genome assemblies using PacBio CCS and PacBio CLR data, we found that our SV benchmark callset is of high quality (Supplementary Table 9). The overall validation rates of insertions and deletions were 95.24% and 95.78% by at least one technology. Although we integrated short-read SV validation callset using 15 SV callers, the validation rates by short-reads (48.7% INS and 76.0% DEL) were much lower than long reads assemblies (90.7% INS and 92.6% DEL). BioNano only validated 3.2% INS and 1.8% DEL over 1 kb, due to its low resolution (kb) by specific restriction enzyme cut sites and failure to accurately determine breakpoints^37^. We also validated our SV benchmark callset with Jia et al.^34^, and found that 97.1% INS and 91.9% DEL were confirmed.

We also compared our SV benchmark calls to the SVs identified by GRC^38^, HGSVC^39^, and HX1^40^ with different groups of samples. The validation rates were 91.3%, 77.8%, and 54.7%, respectively. The high validation rate of GRC was because a Chinese sample was included, and the SVs were also detected from long-read data. Note that such comparison based on limited a number of samples will only detect the common SVs that are shared in different samples.

To defined SV benchmark regions, we used ∼100x PacBio Sequel CLR reads to establish haploid *de novo* assemblies for the parents F7 and M8 (2.94-2.99 Gb), and diploid *de novo* assemblies for the twins D5 and D6 (2.87-2.88 Gb). We then mapped *de novo* assemblies to the GRCh38 reference genome, and ∼2.74∼2.78 Gb callable regions were retained which were supported by reads larger than 50 kb and with mapping quality greater than 5. Regions of assembly-specific SVs, centromeres, and gaps were excluded from callable regions (Supplementary Fig. 7). The Quartet SV benchmark regions cover ∼2.62 Gb of the reference genome (GRCh38; chr1-22) and contains ∼12,705 (75.7%-83.6%) SVs of the benchmark calls. Only SVs inside the benchmark regions are considered when we evaluate variants calling performance of test sets based on benchmark sets with precision and recall.

### Applications of the Chinese Quartet Genomic Reference Materials

#### Evaluating variants calling performance by pedigree information and benchmark sets

We used the whole-genome variants callsets derived from various library preparation methods, sequencing platforms, and bioinformatic tools to demonstrate the usability of the Quartet DNA reference materials in evaluation of variants calling performance. Each callset was evaluated based on the F1-score in the benchmark regions and the Mendelian consistent rate (MCR) on the whole genome.

Four common mappers (Bowtie2, BWA, ISAAC, and Stampy) and six germline variant callers (HaplotypeCaller, ISAAC, Varscan, FreeBayes, Samtools, and SNVer) were compared based on ∼30x Illumina short-read replicates from three sequencing centers (Fig. 4a). Callers had greater impact on variants calling accuracy compared with mappers. SNVs calling performance was high and similar (F1-score 0.978±0.012, MCR 0.944±0.017) among different callers, while Indels calling performance was lower and varied (F1-score 0.732±0.158, MCR 0.695±0.094). RTG, Sentieon, and HaplotypeCaller showed higher F1 scores for indel calling, with Samtools and SNVer performing the worst.

**Figure 4.**
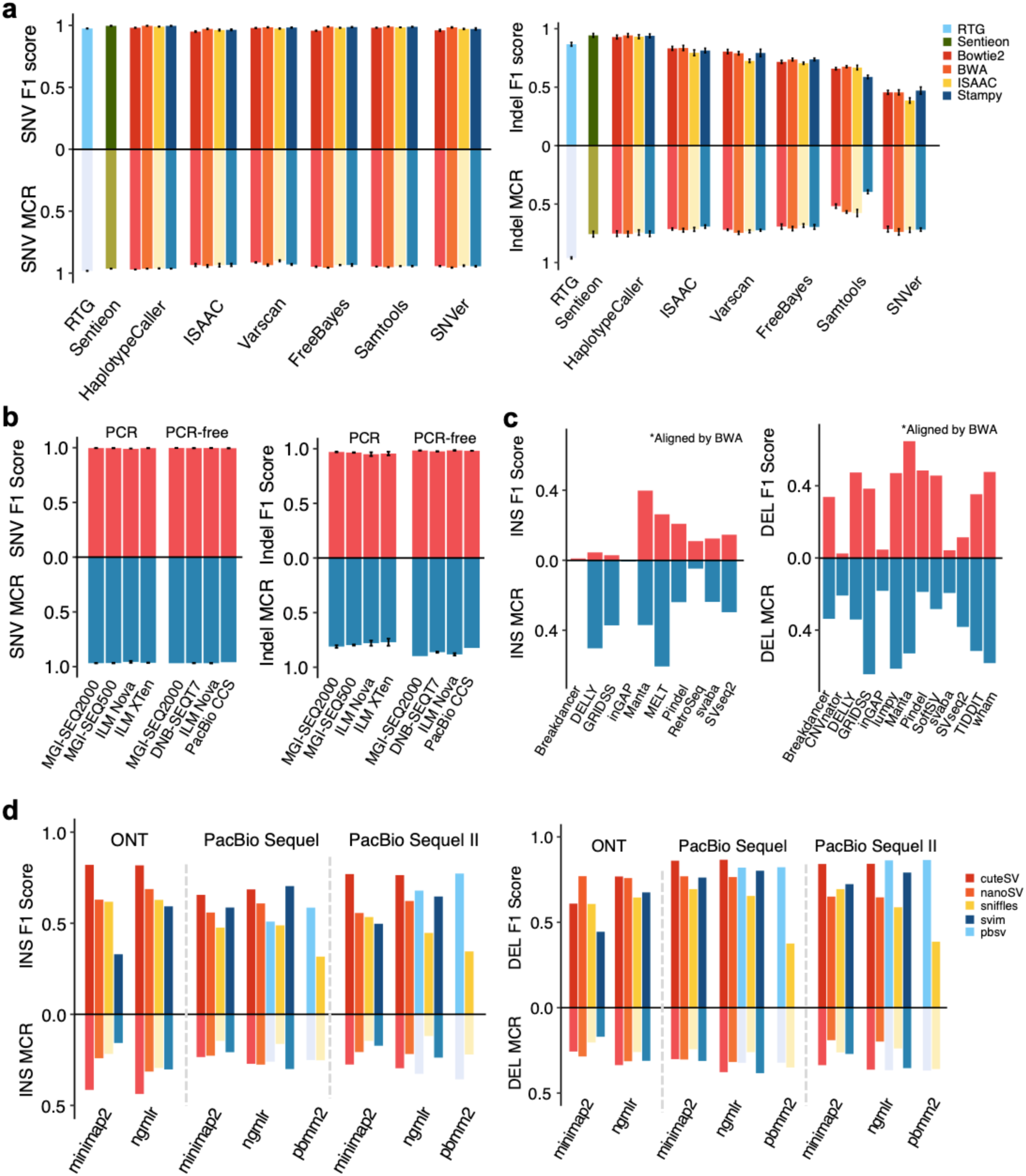
Evaluating variants calling performance by pedigree information and benchmark sets. F1 score and MCR rate of different (**a**) mappers and callers for detecting small variants using Illumina short reads; (**b**) sequencing platforms and library preparation methods for detecting small variants; (**c**) callers for detecting SVs using Illumina short reads; and (**d**) sequencing platforms, combination of mappers and callers for detecting SVs using long-read data.

To investigate the small variants calling performance of different sequencing platforms, we called small variants using the same pipeline (Sentieon) for short-read data and DeepVariant for PacBio CCS reads (Fig. 4b). Illumina platforms, BGI platforms, and PacBio CCS had similar performance, with no obvious differences. Sequencing platforms had smaller impact on variants calling accuracy compared with library preparation methods. PCR-free libraries were superior to PCR libraries for detecting Indels, with higher F1 scores (0.983±0.005 vs 0.958±0.016) and MCR rates (0.921±0.050 vs 0.873±0.094).

For investigating SV calling performance, we compared 15 common callers using short-read data (Fig. 4c). Different callers had various SV calling performance, with F1 scores ranging from 0 to 0.891 and MCR rates ranging from 0 to 0.645. Detection of DEL by short reads was slightly accurate than INS. Only Manta exhibited relatively high F1-score and MCR rate for both INSs and DELs compared to other callers. The MCR of INSs called by DELLY, GRIDSS, and MELT was much higher than F1-score evaluated by benchmark calls, because they detected fewer variants and had lower recall rates. We observed that most callers achieved high performance of DELs results, except for CNVnator, inGAP, and svaba. inGAP identified many more DELs (60,151) than benchmark calls, but had low precision and recall at the same time, indicating its low accuracy.

We also investigated SV calling performance of long-read sequencing platforms and bioinformatic pipelines, by retrospectively evaluating the performance of structural variants call sets used in this study to establish the benchmark sets (Fig. 4d). Generally, more SVs were detected from long reads (7726±3203) than short reads (4922±12471), and present sequencing technologies and algorithms display higher performance for DELs detection than INSs. Combination of mappers and callers should be carefully chosen according to sequencing platforms, since different combinations had F1 scores ranging from 0.374 to 0.856 and MCR rates ranging from 0.119 to 0.437. NGMLR with cuteSV showed high performance detecting both DELs and INSs on all three long-read sequencing platforms. Pbmm2 with pbsv, which was specifically developed for the PacBio platform, performed better on PacBio Sequel II than Sequel. Notably, DELs detected by pbmm2/sniffles had low F1-score but high MCR. Compared with the median het/homo ratio 2.2:1 in 30 call sets, het/homo ratio of pbmm2/sniffles was 0.02:1, which resulted in ∼98% SVs of all four individuals with 1/1 genotypes, indicating that the genotypes of the pipeline were unreliable.

We found that an average of 9% SNVs, 40% indels, 33% DELs, and 20% INSs were located outside the benchmark regions, which could not be evaluated by benchmark sets. The F1 scores for variants inside the benchmark regions might not reflect the accuracy outside the benchmark regions (Supplementary Fig. 8a). As expected, the error rates were significantly higher outside of the benchmark regions. Moreover, the Quartet family design identified more false-positive variant candidates compared to twins and trios and enabled a more precise measurement of error rates (Supplementary Fig. 8b).

#### Identifying and mitigating batch effects in genomic sequencing

To identify batch effects in WGS using the Quartet DNA reference materials, we performed principal component analysis (PCA) on genotype calls detected from various short-read sequencing platforms. Compared with RNA sequencing, DNA sequencing revealed a much smaller level of batch effect^41^. In the scatterplot of the first two eigenvectors, the monozygotic twin daughters were clustered together and located in the middle between the two parents in PC1 and above the parents in PC2, as expected (Supplementary Fig. 9). We observed a clear batch effect from the third and the fourth eigenvectors (Figs. 5a-d). The sequencing platforms played an important role in leading to such detectable batch effects. Large insertions exhibited the lowest reproducibility across the sequencing platforms compared with other variant types, because obvious batch effects were observed even from the first two eigenvectors. Variants called outside the benchmark regions showed larger batch effects than variants called inside the benchmark regions, as expected, because more variants outside the benchmark regions could not reach agreement among call sets (Supplementary Fig. 10).

**Figure 5.**
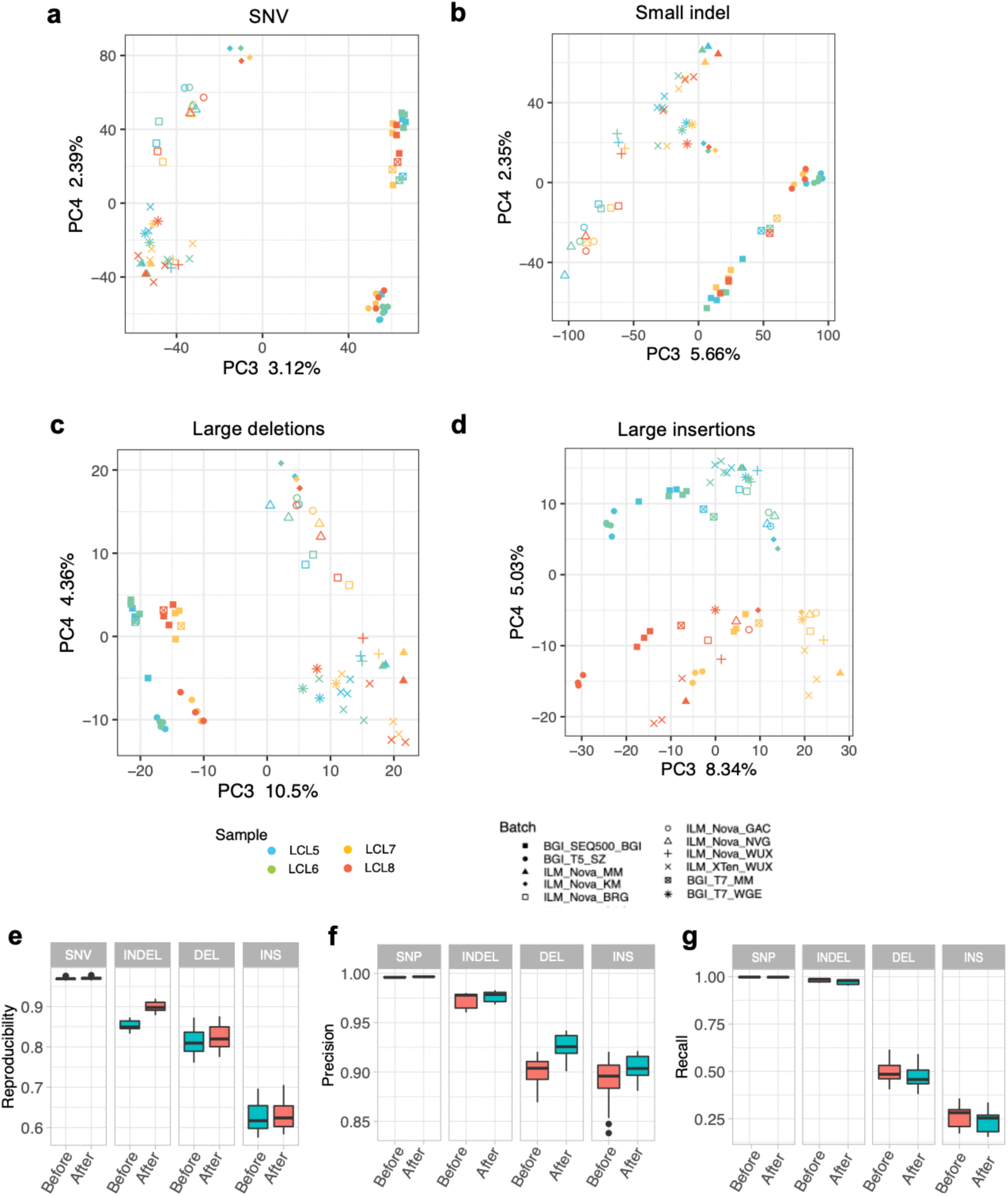
Quartet DNA reference materials can be used to identify and mitigate batch effects in DNA sequencing. The scatterplots of the third and the fourth eigenvectors generated from PCA show batch effects in **(a)** SNVs, **(b)** small indels, **(c)** large deletions, and **(d)** large insertions. **(e)** The percentage of Mendelian concordant and discordant variants as decreasing quality score stringency thresholds were set from left to right. **(f)** Reproducibility of variants called on the whole-genome region before and after filtration. The metrics were defined in our companion paper^33^. **(g)** Precision of variants called inside the benchmark regions before and after filtration. **(h)** Recall of variants called inside the benchmark regions before and after filtration.

Batch effects can be mitigated by removing false positive variants in each batch due to different variant quality metrics such as quality scores, read depth, and mapping quality scores. Pedigree information of the Quartet DNA reference materials can be used to select proper thresholds of those variant quality metrics for each batch to filter potential artifacts. We trained a one-class SVM (support vector machines) classifier using variant quality metrics of Mendelian consistent variants (reliable variants) from one of the three replicates for each batch (Supplementary Table 10, batches 5, 6, and 7). Then the trained models were applied on the other two replicates to filter potential false positives for each batch. The efficiency of batch-specific filtration method was assessed by precision, recall, and cross-batch reproducibility (Figs. 5e-g). After filtration, the cross-batch reproducibility was greatly improved. The precision compared with the benchmark calls increased, while the recall rates decreased, which indicating that false positives were greatly reduced with inevitably sacrificing a small number of true variants.

#### Evaluating variants called from mRNA and protein

Apart from DNA reference materials, we also established RNA, protein, and metabolite reference materials from the same large batch of lymphoblastoid cell lines. Multiomic reference materials from the same resources of Quartet cell lines provide possibilities for cross-validating biological findings from one omics dataset by other levels of omics datasets, supporting quality assessment of a wide range of new technologies and bioinformatics algorithms.

We illustrated a cross-omics validation of variants detected using the Quartet genomics, transcriptomics, and proteomics datasets. As shown in Fig. 6a, a total of 57,000 RNA variants and 18 missense single amino acid variants were detected in RNA-seq and LC-MS/MS based proteomics of Quartet D5, respectively. Specifically, about 40-60% RNA variants in RNA-seq could not be validated by DNA small variant benchmark calls (Fig. 6b). We found more A>G and T>C RNA mutations in the false positive variants than in the true positive variants (Fig. 6c), indicating that RNA editing may play an important part in the high level of inconsistency between variants called from DNA and RNA sequencing datasets. The most prevalent RNA editing involves the A>I conversion (adenosine to inosine), which is recognized by the cellular machineries or sequencing enzymes as A>G (or T>C on the opposite strand) in sequencing data^42^. Fig. 6d shows that a specific SNV benchmark call can be validated by both RNA and protein sequencing data. A missense SNV (chr17:74,866,471 T>C) caused a single amino acid mutation, changing from glutamic acid to arginine.

**Figure 6.**
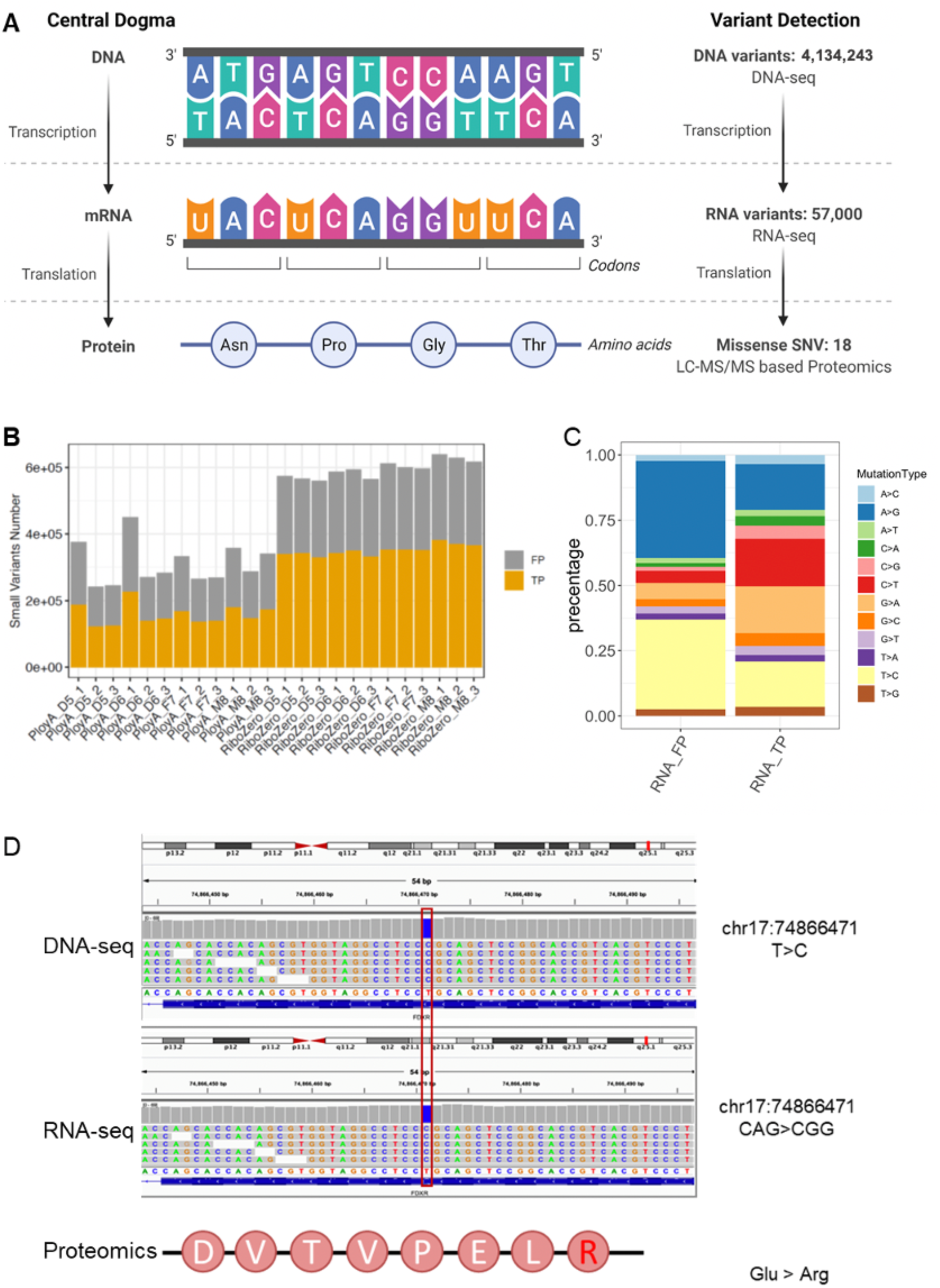
Evaluating variant calling accuracy from RNA and protein data by benchmark calls constructed from DNA data. (**a**) Schematic of central dogma and the number of variants detected in the Quartet DNA-seq, RNA-seq, and LC-MS/MS based proteomics datasets. (**b**) Validation of Quartet RNA variants using DNA reference datasets. True positive (TP) means RNA variants validated in DNA reference datasets, whereas false positive (FP) means the RNA variants not included in the DNA reference datasets. (**c**) Composition of RNA variant types in false positive (RNA_FP) and true positive RNA (RNA_TP) variant calls. (**d**) A T-to-C variant (located in chr17: 74866471) detected by both DNA-seq and RNA-seq is visualized in IGV. The corresponding Glu-to-Arg variant was also detected by LC-MS/MS based proteomics.

These preliminary cross-omics validation results implicated that current applications for variant detection from RNA sequencing and LC-MS/MS based proteomics remain a challenge. The Quartet multiomic reference materials and datasets enable objective quality assessment of these emerging bioinformatics algorithms from cross-omics validations.

## Discussion

One primary challenge of germline variants performance assessment by a single reference sample is that the benchmark sets focus on evaluating the performance of easily detected variants and genomic regions, but ignore difficult variants outside the benchmark regions. Here, we established four DNA reference materials from a Chinese Quartet with parents and monozygotic twins. We constructed high-quality germline benchmark calls, including SNVs, small indels, large insertions and deletions, for each Quartet reference sample based on extensive short-read and long-read sequencing. The quality of the benchmark calls was improved through a series of data-filtering procedures including consensus voting of replicates, pedigree information, and orthogonal technologies.

We demonstrated that the use of the Quartet DNA reference materials together helps make a comprehensive performance assessment of variants across the whole genome. There are two aspects of “truth” related to the Quartet DNA reference materials. One aspect is related to the benchmark calls, where only highly confident variants were kept. Precision and recall are commonly used metrics to evaluate variants calling performance within benchmark regions. Another aspect is the genetic built-in truth of the monozygotic twins and their parents. Mendelian concordance rate of variants among the Quartet members can be used to estimate the fraction of variants that might be correctly detected. Compared to other studies focusing on easy-to-detect variants in benchmark regions alone, difficult variants outside the benchmark regions not only reflect major discordances among different sequencing platforms and labs, but also help guide future development and optimization of sequencing technologies.

There are also drawbacks only using pedigree information instead of benchmark sets for performance assessment. For example, systematic sequencing or mapping errors, such as heterozygotic or homozygotic variants called on all Quartet samples, which are Mendelian consistent, will be mistakenly considered as true variants. In some cases, Mendelian concordance rate is low due to sequencing failure of one or more Quartet reference samples. Comparison with the benchmark calls can help identify which sample exhibits bad variants calling performance. Notably, pedigree information can be used to evaluate Mendelian concordance, but it cannot help determine false negatives. Therefore, benchmark sets are necessary to identify false negatives and measure recall rate, while the pedigree information provides additional tool for the assessment of variants calling accuracy outside the benchmark regions.

Although benchmark sets are important, the current version of benchmark regions are relatively limited. Future work is needed to combine short-read and long-read data to expand the benchmark regions by resolving more difficult genomic regions. We will also include inversions, duplications and translocations in the benchmark sets as methods for variant detection improve.

To evaluate and monitor the performance of the data generation processes, sequencing all the Quartet genomes is not cheap, especially for long-read sequencing. If one is only interested in variants or regions in the benchmark calls and regions, we recommend sequencing one of the Quartet samples and making quality assessment using benchmark sets by precision and recall. If the aim is to improve current technologies in some challenging genomic regions, we recommend sequencing all four Quartet samples to estimate performance on those difficult regions. Since a new technology is often accompanied by advantages beyond what current technologies can offer, the Quartet based Mendelian concordance rate is independent of the benchmark calls and can provide a more objective evaluation.

To monitor and improve data quality across different sequencing centers in large-scale studies, we recommend sequencing all the Quartet DNA reference materials per batch. In an automated library preparation setup, 96 samples are routinely handled in a batch. Although including four quality control samples per batch increases experimental cost by ∼5%, it can benefit the study tremendously by identifying and mitigating batch effects for the sake of discovering genuine biomarkers for precision medicine.

The Quartet DNA and other types of omics reference materials are publicly available to the community by requesting through the Quartet Data Portal website (http://chinese-quartet.org/). We encourage researchers to upload and share Quartet sequencing data, thereby hoping the rich collections of diverse datasets and analysis for the Quartet samples will enable optimization of the benchmark sets and regions. In summary, the Quartet DNA reference materials and datasets are essential resources for objective and comprehensive evaluation of the quality of sequencing and bioinformatic methods, which will greatly improve the quality control awareness of the sequencing community and help overcome barriers to the translation of findings from genomic studies into clinical practices.

## Methods

### Materials Availability

The Quartet DNA reference materials generated in this study can be requested from the Quartet Data Portal (http://chinese-quartet.org/) under the Administrative Regulations of the People’s Republic of China on Human Genetic Resources.

### Data Availability

Short-read and long-read datasets used in this study can be obtained through the Genome Sequence Archive GSA (GSA) of the National Genomics Data Center of China with BioProject ID of PRJCA007703, or from SRA. Quartet DNA benchmark sets generated by this study can be accessed from the Quartet Data Portal (http://chinese-quartet.org/).

### Establishing DNA reference materials

The Chinese Quartet DNA reference materials were extracted from four immortalized B-lymphoblastoid cell lines transfected by Epstein-Barr virus, including father (F7), mother (M8) and monozygotic twin daughters (D5/D6). We extracted two batches of DNA on August 6, 2016 and October 28, 2017 from two large expansions of the cell lines. We diluted DNA to 220 ng/μL and made >1,000 aliquots for each DNA sample. Each vial contains 10 μg of DNA in TE buffer (10 mM TRIS, pH 8.0; 1 mM EDTA, pH 8.0). The Quartet DNA is stored at −80°C for long-term preservation, or at 4°C for short-term preservation. We checked the integrity of DNA (DIN) by Agilent 4200 and the distribution of DNA fragment length by Agilent 2200. The Quartet DNA is stable for at least three years at −80°C and for three weeks at 4°C during the entire duration of quality monitoring. This study focuses on germline variants calling quality control. Two batches (Lot 20160806 and Lot 20171028) of DNA reference materials were extracted from large expansion of cell lines, with 1000 tubes (10 μg, 220 ng/μL) for each Quartet reference sample at each batch. DNA reference materials are stable and in good quality. The peak size of DNA fragments is over 60 kb. The stability has been monitored monthly for three years, with DNA integrity number (DIN) over 8.5.

### Library preparation and whole-genome sequencing

#### 1. Short-read sequencing

Twelve tubes of Quartet DNA reference materials, with three replicates for each of the four Quartet sample types, were sequenced per batch. DNA reference materials were from Lot 20160806. We obtained datasets from four sequencing platforms in six sequencing labs by PCR and PCR-free library protocols, resulting in 27 replicates per sample and 108 libraries in total:

1. ∼50x paired-end, whole-genome sequencing with 2×100 bp reads of ∼250 bp insert size from MGI-SEQ2000 with PCR library kit, performed at BGI.
2. ∼30x paired-end, whole-genome sequencing with 2×150 bp reads of ∼300 bp insert size from Illumina HiSeq XTen with TruSeq Nano library kit, performed at ARD and NVG.
3. ∼30x paired-end, whole-genome sequencing with 2×150 bp reads of ∼400 bp insert size from Illumina HiSeq XTen with TruSeq Nano library kit, performed at WUX.
4. ∼30-60x paired-end, whole-genome sequencing with 2×150 bp reads of ∼300-400 bp insert size from Illumina NovaSeq6000 with PCR-free library kit, sequenced at ARD, BRG, and WUX.
5. ∼35x paired-end, whole-genome sequencing with 2×150 bp reads of ∼380 bp insert size from DNB-SEQT7 with PCR-free library kit.

#### 2. Long-read sequencing

To establish structural variant benchmark calls, the four Quartet DNA reference materials, one replicate for each sample, were sequenced on three long-read platforms, resulting in three libraries per sample and 12 libraries in total:

1. ∼100x, whole-genome sequencing with 11-14 kb mean read length and 20-25 kb N50 read length from Oxford Nanopore Technologies (ONT). DNA reference materials were from Lot 20171028.
2. ∼100x, whole-genome sequencing with 8-11 kb mean read length and 13-18 kb N50 read length from PacBio Sequel (CLR). DNA reference materials were from Lot 20160806.
3. ∼30x, whole-genome sequencing with 16-18 kb mean read length and 26-28 kb N50 read length from PacBio Sequel II (CLR). DNA reference materials were from Lot 20160806.

We also generated sequencing datasets from BioNano, 10x Genomics and PacBio CCS reads to validate benchmark calls:

1. BioNano Genomics: ∼200X for D5, ∼300X for D6, F7 and M8 BioNano Genomics data with average fragment length 260∼300 kb. DNA reference materials were from Lot 20160806.
2. 10x Genomics: ∼30X Genomics data with average fragment length ∼150 kb. DNA reference materials were from Lot 20160806.
3. ∼50x, whole-genome sequencing with 13-14 kb mean read length and 13-14 kb N50 read length from PacBio Sequel II (CCS HiFi reads). DNA reference materials were from Lot 20160806.

### Reads mapping and variants calling for short-read sequencing

Sequences were mapped to GRCh38 (https://gdc.cancer.gov/about-data/gdc-data-processing/gdc-reference-files). We used Sentieon Genomics software (https://www.sentieon.com/) to analyze short-read WGS datasets from raw fastq files to GVCF files. This workflow was derived from recommended germline small variants calling pipeline by the Broad Institute (https://gatk.broadinstitute.org/hc/en-us/articles/360035535932-Germline-short-variant-discovery-SNPs-Indels-), including reads mapping by BWA-MEM, duplicates removing, indel realignments, base quality score recalibration (BQSR), and variants calling by HaplotyperCaller in GVCF mode. Then we performed joint variants calling using Sentieon GVCFtyper to merge all 108 GVCF files. We used default settings for all processes.

Different from regular VCFs, GVCF files have records and extra information for all genomic sites. A site is recorded as a variant call, homozygotic reference, or with no reads covered. In a regular VCF, we cannot distinguish a site with no information from a homozygotic reference. GVCF files enable us avoid mistaking no-call sites as homozygotic references, and facilitate representation of complex variants as well.

To keep as many variants as possible and not to remove any potential true variants with low qualities, we did not filter variants from the original GVCF call sets by empirical variants quality or machine learning based variants quality score recalibration (VQSR).

### Reads mapping and variants calling for long-read sequencing

We used three mappers (NGMLR, minimap2, and pbmm2) and five callers (cuteSV, NanoSV, Sniffles, pbsv, and SVIM) to call structural variants, resulting in 11 combinations. Reads were mapped to human genome version hg38 (GCA 000001405.15) from UCSC Genome Brower (http://hgdownload.soe.ucsc.edu/goldenPath/hg38/chromosomes/).

PacBio Sequel-based call sets were generated as follows:

1. Reads were aligned with NGMLR v.0.2.7 with -x pacbio parameter, minimap2 v.2.17-r941 with -x map-pb --MD -Y parameters and pbmm2 v.1.0.0 with --sort --median-filter --sample parameters separately.
2. Structural variants calling was performed using cuteSV v.1.0.4 with --genotype parameter, NanoSV v1.2.4 with per chromosome pattern and an ancillary file containing random positions in hg38, Sniffles v.1.0.11 with default parameter and SVIM v.1.2.0 with --minimum_depth 10 parameter based on BAM files created by NGMLR v.0.2.7 and minimap2 v.2.17-r941 separately. Additionally, Sniffles v.1.0.11 was also run on pbmm2 v.1.0.0 and pbsv v.2.2.1 was run on pbmm2 v.1.0.0 and NGMLR v.0.2.7. The pbsv discover stage was run with --tandem-repeats parameter using tandem repeat annotations file human_GRCh38_no_alt_analysis_set.trf.bed (https://github.com/PacificBiosciences/pbsv/tree/master/annotations). The pbsv discover and call stages were both run on the full genome.

PacBio Sequel II-based call sets were generated as follows:

1. Reads were aligned with NGMLR v.0.2.7 with -x pacbio parameter, minimap2 v.2.17-r941 with -x map-pb --MD -Y parameters and pbmm2 v.1.0.0 with --sort --median-filter --sample parameters separately.
2. Structural variants calling was performed using cuteSV v.1.0.4 with -s 3 --genotype parameters, NanoSV v1.2.4 with per chromosome pattern and an ancillary file containing random positions in hg38, Sniffles v.1.0.11 with -s 3 parameter and SVIM v.1.2.0 with --minimum_depth 3 parameter based on BAM files created by NGMLR v.0.2.7 and minimap2 v.2.17-r941 separately. Additionally, Sniffles v.1.0.11 with -s 3 was also run on pbmm2 v.1.0.0 and pbsv v.2.2.1 was run on pbmm2 v.1.0.0 and NGMLR v.0.2.7. The pbsv discover stage was consistent with PacBio Sequal-based process.

Nanopore-based call sets were generated as follows:

1. Reads were aligned with NGMLR v.0.2.7 with -x ont parameter and minimap2 v.2.17-r941 with -x map-ont --MD -Y parameters.
2. SVs were called using cuteSV v.1.0.4 with --genotype parameter, NanoSV v1.2.4 with per chromosome pattern and an ancillary file containing random positions in hg38, Sniffles v.1.0.11 with default parameter and SVIM v.1.2.0 with --minimum_depth 10 parameter based on BAM files created by NGMLR v.0.2.7 and minimap2 v.2.17-r941 separately.

In addition to the parameters of mappers and callers mentioned above, the others are default.

### Detecting structural variants from Illumina-based short-read sequencing

Illumina NovaSeq WGS short-read sequencing with ∼40x 2×150 bp and 420 bp insert size was performed at ARD and used to call structural variants. The reads were mapped to the GRCh38.d1.vd1 reference genome by Sentieon BWA. According to previous studies^43^, 15 algorithms with relatively high precision and/or recall were selected for structural variants discovery, including Breakdancer^44^, CNVnator^45^, DELLY^46^, GRIDSS^47^, inGAP-sv^48^, LUMPY^49^, Manta^50^, MELT^51^, Pindel^52^, softSV^53^, SvABA^54^, SVseq2^55^, tardis^56^, TIDDIT^57^, and Wham^58^. Consequently, 15 Illumina-based call sets were generated for each Quartet reference sample. Structural variants were filtered based on the number of reads supporting structural variants (RSS), types and lengths. For several algorithms, RSS value was not available and other values such as quality scores were used to simulate RSS. Only five types of structural variants were retained (INSs, DELs, DUPs, INVs, and BNDs). Structural variants under 50 bp were removed except for BNDs. The filtered output file for each algorithm was converted to a VCF format with SVMETHOD, END, SVTYPE, and SVLEN tags in the information field. All 15 call sets for each individual in Quartet were merged into a single call set based on the same type and with breakpoints distance of 1 kb using SURVIVOR v.1.0.7.

### Detecting small variants and structural variants from 10x Genomics linked reads

Small variants and structural variants were called by longranger-2.2.2 (https://support.10xgenomics.com/genome-exome/software/downloads/latest) with default parameters from 10x Genomics linked reads data sets. Small variants were from phased_variants.vcf.gz. Structural variants over 50bp were from dels.vcf.gz and large_svs.vcf.gz were retained.

### Detecting structural variants from BioNano

The structural variants were called by BioNano Solve v3.1 (bnxinstall.com/solve/Solve3.1_08232017) with default parameters.

### Detecting small variants and structural variants from PacBio CCS reads

The small variants were called by DeepVariant (https://github.com/google/deepvariant) with default parameters. The structural variants were called by pbsv (https://github.com/PacificBiosciences/pbsv) with default parameters.

### Detecting structural variants from PacBio assembly alignments

The complete diploid assembly was reconstructed based on trio binning of canu v1.8 from PacBio Sequel CLR data (∼100X) of twins and Illumina NovaSeq ∼40x 2×150bp WGS short-read sequencing data with 420 bp insert size performed at ARD for the two parents. Trios are formed by twins D5 and D6 and their parents respectively. Each trio is then assembled independently. The assembly was performed with canu -p prefix -d prefix genomeSize=3.1g -pacbio-raw pacbio.fasta.gz -haplotypeF7 F7.NGS.fastq.gz -haplotypeM8 M8.NGS.fastq.gz. The diploid assembly results of parents were generated by FALCON v0.4 with default parameters based on ∼100x PacBio Sequel CLR sequencing data.

Two methods of assembly alignment were used, including MUMmer v4.0.0beta2 and minimap2 v.2.17-r941. MUMmer assembly alignments were performed with the commands nucmer -maxmatch -l 100 -c 500 ref.fa --prefix haplotype.contigs.fasta. Minimap2 assembly alignments were performed with the commands minimap2 -cx asm5 -t12 --cs ref.fa haplotype.contigs.fasta. Three assembly-based callers were used including Assemblytics V1.2.1, SVMU V0.4, and Paftools (https://github.com/lh3/minimap2/tree/master/misc). Assemblytics was run with the parameters unique_length_required=10000 min_size=20 max_size=1000000 by MUMmer alignment. Results were transformed into VCF format using SURVIVOR. SVMU was run with default parameters by MUMmer alignment. Paftools was run with default parameters to identify structural variants from the CS tags generated by Minimap2 alignment. Results of SVMU and Paftools were transformed into VCF format using a custom script. Structural variants of two contigs of the twins were merged into a single call set, and then structural variants shared between twins are used to validate structural variants benchmark calls.

### Preprocessing and filtering of structural variants call set from long-read sequencing

Due to considerable diversity in the number, type, and size of structural variants and the format of VCF files created by different caller algorithms, it was difficult to merge the original VCF files directly for downstream analysis. In order to unify the standard and facilitate the analysis, structural variants call sets were preprocessed as follows:

1. Only five types of structural variants (INS, DEL, DUP, INV, and BND) were retained for each call set. For Sniffles, complex structural variant types were excluded. For SVIM, DUP_INT, and DUP:TANDEM were converted to DUPs. For pbsv, CNVs were filtered.
2. All structural variants under 50 bp were removed except for BNDs.
3. All structural variants call sets were filtered if they do not meet the minimum number of supporting reads. For ∼100X PacBio Sequel and ONT sequencing datasets, structural variants call sets from cuteSV and Sniffles were filtered with tag RE ≥10. SVIMs were filtered with tag SUPPORT ≥10. Structural Variants called by NanoSV were filtered with tag DV ≥10. For pbsv, structural variants were filtered based on read depth of variant allele ≥3 of tag AD. The parameter median-filter in pbmm2 v.1.0.0 only aligns the subread closest to the median subread length per ZMW and significantly reduces the number of reads supporting structural variants, thus a lower filtering threshold should be used. Otherwise, pbsv will lose too many true variants. For ∼30X PacBio Sequal II, heuristically, the minimum number of reads supporting structural variants in all call sets from cuteSV, Sniffles, SVIM, and NanoSV was adjusted to three. The filtering threshold of pbsv was the same as that of PacBio Sequel for the parameter --median-filter.
4. All structural variants call sets were assigned a unique ID based on sequencing platform, sample name, pipelines, serial number and structural variants type for backtracking easily.

### Integration of small variant benchmark calls

Construction of small variants benchmark calls was described as follows:

1. GVCF files of 108 libraries were merged by joint variants calling process for each chromosome separately (chr1-22, X), with samples in columns and variants in rows.
2. Variants recurrent in most of 27 replicates for each Quartet sample were selected. This process was run for each chromosome. Then we merged results of all chromosomes and obtained four integrated catalogs corresponding to the four Quartet reference samples. Each site was annotated by “Conflict” (fail in the replicate consensus process), “./.” (no call in most replicates), or agreed genotype by voting.
3. We removed sites annotated as “Conflict” in any of four Quartet samples, and sites voted as “0/0” or “./.” in all four Quartet samples.
4. A total of 31,155 small variant positions overlapping deletions were removed in all four Quartet samples, which represented with “*” in gvcf files, because downstream analysis tools cannot del with * allele.
5. Mendelian inheritance status of remaining sites was checked by VBT ^32^. We split Quartet into two “trios” (D5-F7-M8 and D6-F7-M8), and performed Mendelian analysis by VBT separately. Only variants shared between twins and Mendelian consistent with parents were retained.
6. We kept variants in callable regions described below for all four Quartet samples.

### Integration of structural variant benchmark calls

The benchmark structural variants were constructed based on all 120 long-read sequencing structural variants call sets described above, only including chr1-22:

1. Structural variants callers with different detection algorithms lead to the same variant being called with different breakpoints and lengths. Moreover, due to the scoring systems of aligners and different clustering methods of callers, some large structural variants events were split into several smaller INSs/DELs in a local region. These redundant variants inflated the number of structural variants and hindered subsequent merging calls between different callers for the same sample. Jasmine v.1.0.1 (https://github.com/mkirsche/Jasmine) uses an improved minimum spanning forest algorithm to merge different variants within a single caller or between callers. Each variant was represented by a breakpoint (start, length) in two-dimensional space. The distance between the two variants was equal to the Euclidean distance (default) by their breakpoints. When the distance between variant breakpoints met the max_dist value (default value 1000, Euclidean distance: [(start1-start2) ^2+ (length1-length2) ^2]1/2 ≤1000), these close variants with the same variant type were clustered into a single structural variant event.
2. We used Jasmine with --allow_intrasample, --keep_var_ids and --ignore_strand parameters to merge structural variants between callsets for each sample.
3. The integrated structural variants set of each individual sample was subsequently filtered to retain structural variants supported by at least two long-read sequencing platforms or at least six call sets in a single technology.
4. The four integrated structural variants sets in Quartet were merged into one call set by Jasmine with --keep_var_ids and --ignore_strand parameters.
5. Structural variants were excluded if their size is over 10 Mb and in low-confidence regions, including centromeres, pericentromeric region and gaps in hg38 reference genome.
6. Structural variants frequently occur on repeats, which seriously hinders accuracy of detecting breakpoints and sequences on the alternative allele. Structural variants with explicit sequences were also helpful for subsequent genotyping. Therefore, we used Iris v1.0.1 to report alternative allele sequences of INSs and DELs. It extracted breakpoints by racon or falcon_sense to get consensus sequences. Then NGMLR or minimap2 was used to re-align these sequences of the breakpoints to the reference genome for refining the variant breakpoints and sequences. The read names of supporting structural variants and allele sequences were obtained by Sniffles with -n -1 -s 2 --Ivcf parameters. We refined INSs and DELs by Iris with max_out_length=1000000, --also_deletions and --pacbio parameters. In addition, the minimap2 bam files from PacBio Sequel II of each Quartet sample were adopted for reporting sequence and refining breakpoints, because PacBio Sequel II sequencing datasets had lower mismatch rates.
7. We re-genotyped merged structural variants from two long-read sequencing platforms (PacBio Sequel and ONT) by three long-read genotypers (LRcaller v0.1.2, Sniffles v1.0.11 and SVJedi v1.1.0) with default parameters. The bam files from NGMLR and minimap2 of PacBio Sequel and ONT were used by Sniffles and LRcaller. The fasta files of PacBio Sequel and ONT were used by SVJdei. Thus, for each Quartet sample, a total of 10 genotyping call sets were produced, four from LRcaller, four from Sniffles and two from SVJedi. SVs were considered successfully and concordantly genotyped if at least six of the ten genotypes were the same.
8. The structural variants successfully genotyped as heterozygous variants or homozygous variants in at least one of four Quartet samples were retained as input of Mendelian analysis. We retained structural variants that were shared by twin daughters and Mendelian consistent with parents, using bcftools v.1.9-224-g96ef00a.

### Defining benchmark regions of small variants

First, we obtained callable regions from bam files using GATK V3.8-1 CallableLoci for each of the 108 short-read libraries, with –maxDepth 300 --maxFractionOfReadsWithLowMAPQ 0.1 -- maxLowMAPQ 1 --minBaseQuality 20 --minMappingQuality 20 --minDepth 10 -- minDepthForLowMAPQ 10 parameters.

We next selected consensus callable bed regions for each Quartet reference sample, if bed regions were denoted as callable (1) at least 2/3 replicates in one batch, (2) at least 4/5 batches by PCR library preparation and 3/4 batches by PCR-free, and (3) both PCR and PCR-free library preparation methods. Then we kept regions callable in all Quartet samples.

We obtained reproducible invariant genomic positions by the same voting process. We then converted reproducible invariant genomic positions and high-confidence small variant positions to bed region, and kept regions where all Quartet samples had concordant voting results.

Benchmark regions include positions of small variants benchmark calls and invariant homozygotic reference positions in consensus callable regions mention above. Thus, we got regions which were callable and had consistent calling results among replicates and all Quartet samples.

### Defining benchmark regions of structural variants

When evaluating analysis methods using structural variants benchmark calls, structural variants were limited in the benchmark regions, which could assess the accuracy of genotyping about INSs and DELs.

The process for constructing benchmark regions was as follows:

1. We first identified callable regions covered by exactly one contig from output of Paftools based on trio binning genome assembly of Canu, as described in PacBio assembly-based structural variants detection. By default, Paftools used assembly-to-reference alignment longer than 10 kb to generate callable regions.
2. For each individual in twins, we got the union of the regions from each parental haplotype. Then we obtained the intersection of callable regions between twins.
3. We compared the benchmark calls and PacBio-based assembly structural variants from Paftools in twins through Jasmine with --keep_var_ids and --ignore_strand parameters, and then retained assembly specific structural variants.
4. We applied svanalyzer widen command to extend the repetitive genomic coordinates surrounding assembly specific structural variants, and then added 50 bp on each side of these regions.
5. Based on the regions obtained in step 2, we removed the regions in the step 4. Finally, we constructed the benchmark regions for benchmark set in twins.
6. The process for constructing benchmark regions in parent was similar to that of twins except for step 2, because there were no biological replicates of the parents.

### Validation of small variants benchmark calls by PMRA

We performed 16 replicates for each Quartet reference material on the Applied Biosystem^TM^ Axiom^TM^ Precision Medicine Research Array (PMRA). Genotypes were called by Axiom Analysis Suite v4.0.1.

We selected genotype calls using the following criteria: (1) less than two replicates with missing calls; and (2) more than 80% genotype calls are the same.

The PMRA probes were annotated by hg19, but the reference datasets were mapped based on GRCh38. To avoid converting errors, we only compared variants annotated in dbSNP by dbSNP RefSNP ID.

### Validation of structural variant benchmark calls by independent technologies

We validated the structural variants benchmark calls using four Illumina short-reads, 10× Genomics linked reads, PacBio CLR long reads and BioNano Genomics optical mapping. The structural variants datasets corresponding to each technology were generated through the data generation section above. In each technology, the shared structural variants between twins were used for validation of benchmark call structural variants in twins. The structural variants benchmark calls in parents were separately validated by the corresponding structural variants datasets. The validation process used Jasmine with --keep_var_ids and –ignore strand parameters.

We also randomly selected 40 structural variants including 20 insertions and 20 deletions that have not been validated by other technologies and manually checked their accuracy through IGV. In addition, the datasets from three other independent researches based on long read sequencing were employed to validate our benchmark calls using Jasmine with --keep_var_ids and –ignore strand parameters.

### Training batch-specific machine learning models

Variant quality metrics of Mendelian concordant variants from one D5 replicate for each batch (Supplementary Table 10, Batches 5, 6, and 7 with three replicates) were used to train one-class SVM classifier (https://scikit-learn.org/stable/modules/svm.html#). For small variants, variant quality, depth, BaseQRankSum, QualByDepth, FisherStrand, SrandOddsRatio, RMSMappingQuality, MappingQualityRnakSunTest, ReadPosRankSumTestg, genotype quality and membership of dbSNP were used. For structural variants, variant quality, genotype quality and the raw counts of paired reads supporting alternate allele were used. The three trained models were applied for each batch respectively to classify high-quality variants and low-quality variants.

### Variant calling from RNA-seq

Sequences were mapped to GRCh38. We used Sentieon Genomics software to analyze short-read RNA datasets from raw fastq files to VCF files. This workflow was derived from recommended RNA-seq short variants discovery pipeline by the Broad Institute (https://gatk.broadinstitute.org/hc/en-us/articles/360035531192-RNAseq-short-variant-discovery-SNPs-Indels-), including reads mapping by BWA-MEM, duplicates removing, split reads at junction, base quality score recalibration (BQSR), and variants calling by HaplotyperCaller. Variants were filtered by GATK VariantFiltration with parameters: -window 35 -cluster 3 -filterName FS -filter “FS > 30.0” -filterName QD -filter “QD < 2.0”.

### Variant detection from LC-MS/MS proteomics

XML file contained peptide identification results generated by an open-source search engine X!Tandem. The software needs to input the Mascot Generic Format (MGF) file, which is the most common format for MS/MS data encoding in the form of a peak list. Then PGA R packages (v1.18.1) were used to identify variant peptides from the XML file.

We constructed custom protein databases from RNA-seq datasets containing SNVs and Indels, then searched the database to detect variant peptides and their corresponding variants locations on the genome from LC-MS/MS datasets.

### Precision and recall

Precision is the fraction of called variants in the test dataset that are true, and recall is the fraction of true variants are called in the test dataset. True Positives (TP) are true variants detected in the test dataset. False Negatives (FN) are variants in the reference dataset failed to be detected in the test dataset. False Positive (FP) are variants called in the test dataset but not included in the reference dataset. Precision and recall are defined as below:

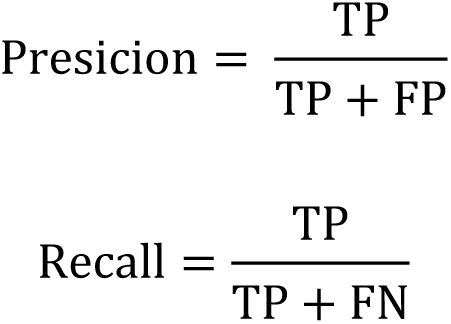

For small variants, we compared variants with benchmark small variants using hap.py (https://github.com/Illumina/hap.py). For structural variants, we merged and compared variants in different callsets using Jasmine with parameters max_dist=1000 --keep_var_ids --ignore_strand. When considering the genotype of structural variants, an additional parameter --output_genotypes needs to be used. When comparing with small variants benchmark calls and structural variants benchmark calls, genotypes of the variants were considered.

### Mendelian violation rate of Quartet family

Mendelian violation rate is the number of variants not following Mendelian inheritance laws divided by the total number of variants called among the four Quartet samples. Mendelian violated variants are the variants not shared by the twins or following Mendelian inheritance laws with parents. When calculating Mendelian violation of small variants, variants on large deletions defined by structural variants benchmark calls were not included, because VBT (https://github.com/sbg/VBT-TrioAnalysis) takes these true variants as Mendelian violations. For structural variants, Mendelian analysis was only done for Quartet-D5, because we could not distinguish homozygotic references and no call sites. We did not consider genotype information; therefore, Mendelian discordant variants are variants not shared by Quartet twins or specifically identified in twins but not in parents.

## Disclaimer

This article reflects the views of the authors and does not necessarily reflect those of the US Food and Drug Administration.

## Acknowledgements

This study was supported in part by National Key R&D Project of China (2018YFE0201603 and 2018YFE0201600), the National Natural Science Foundation of China (31720103909 and 32170657), Shanghai Municipal Science and Technology Major Project (2017SHZDZX01), State Key Laboratory of Genetic Engineering (SKLGE-2117), and the 111 Project (B13016). Some of the illustrations in this paper were created with BioRender.com.

## Author contributions

Y.Z., X.F., S.F., J.W., H.H., and L.S. conceived and oversaw the study. Y.Z., L.D., R.Z., and W.H. prepared the biosamples and coordinated NGS library preparation and sequencing. L.R., X.D., J.Y., Y.G., R.P., Y.L., J.L., Y.Y., N.Z., J.S., F.L, D.W, H.C., L.L.S., L.H., A.S., J.N., W.X., J.X., W.T., X.H., P.J., K,Y., J.M.L., L.J., L.S., H.H., J.W., S.F., X.F., and Y.Z. performed data analysis and interpretation. J.Y., L.R., Y.L., and J.S. managed the datasets. R.Z., L.R., R.P., J.M.L, L.S., and Y.Z. coordinated proficiency testing with the Quartet DNA reference materials. L.R., X.D., Y.Z., S.F., H.H., L.S., W.T., J.X., and W.X. wrote and revised the manuscript. All authors reviewed and approved the manuscript. Dozens of participants of the Quartet Project freely donated their time and reagents for the completion and analysis of the project.

## Supplementary Figures

**Supplementary Figure 1.**
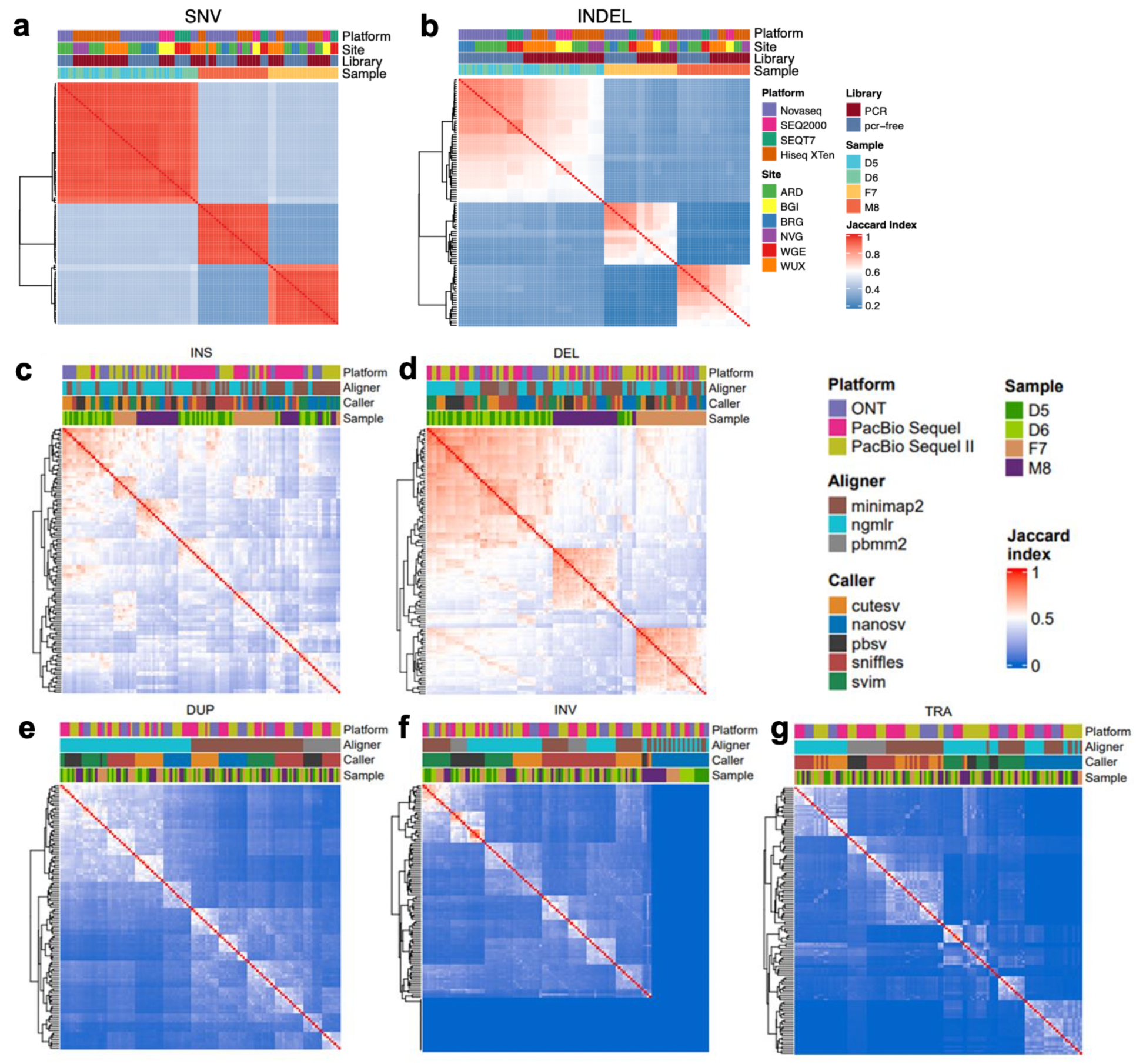
Pairwise comparison of (**a**) SNVs, (**b**) small indels (**c**) insertions, (**d**) deletions, (**e**) duplications, (**f**) inversions, and (**g**) translocations called from short-read and long-read WGS call sets used to establish benchmark calls. Color of each cell corresponds to Jaccard index (the number of shared variants divided by the union of two call sets), with high similarity in red and low similarity in blue.

**Supplementary Figure 2.**
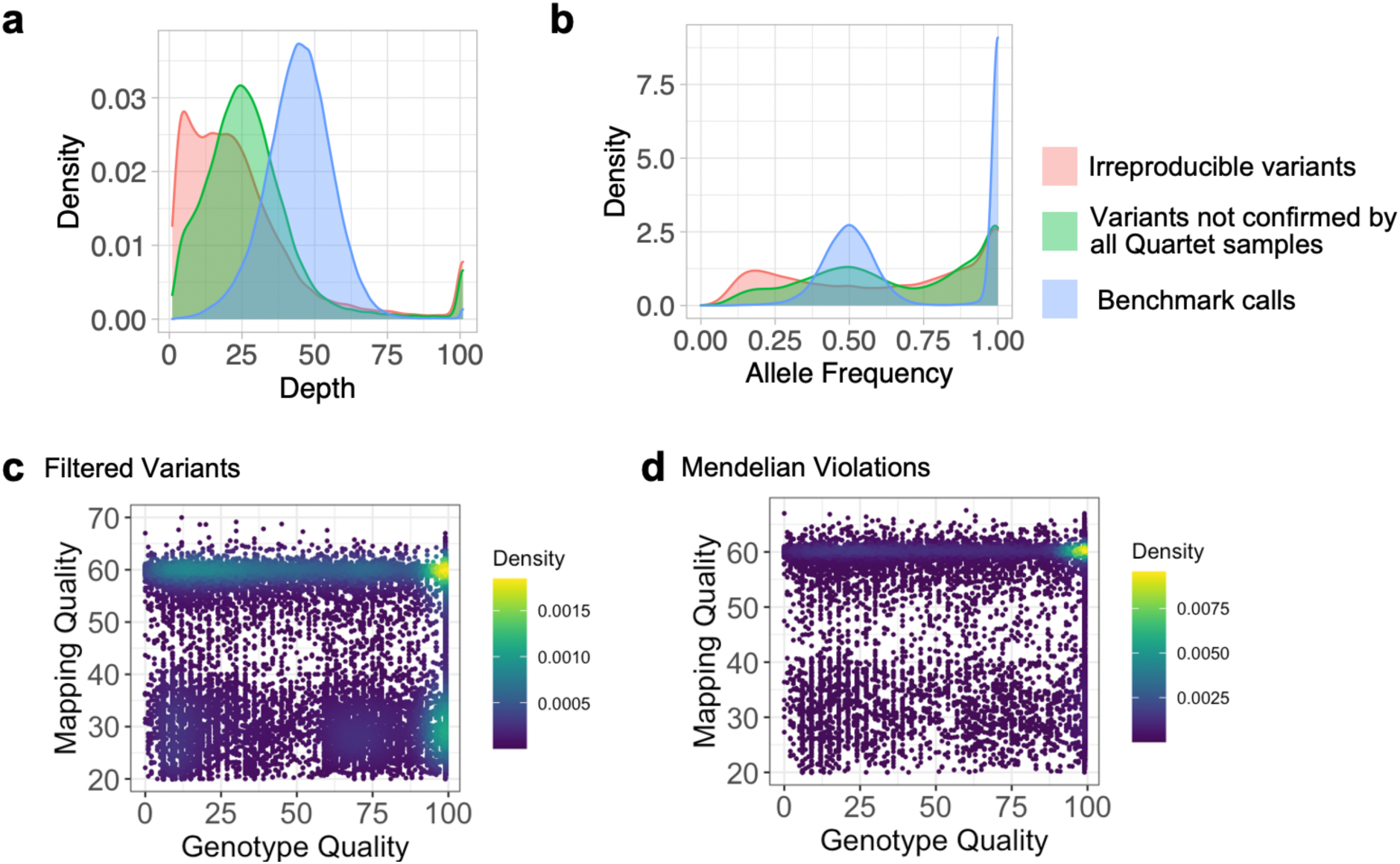
Density plots show differences in (**a**) sequencing depth, (**b**) allele frequency, (**c** and **d**) genotype quality and mapping quality between inconsistent small variants across call sets, consistent Mendelian discordant small variants, and reproducible and Mendelian concordant small variants.

**Supplementary Figure 3.**
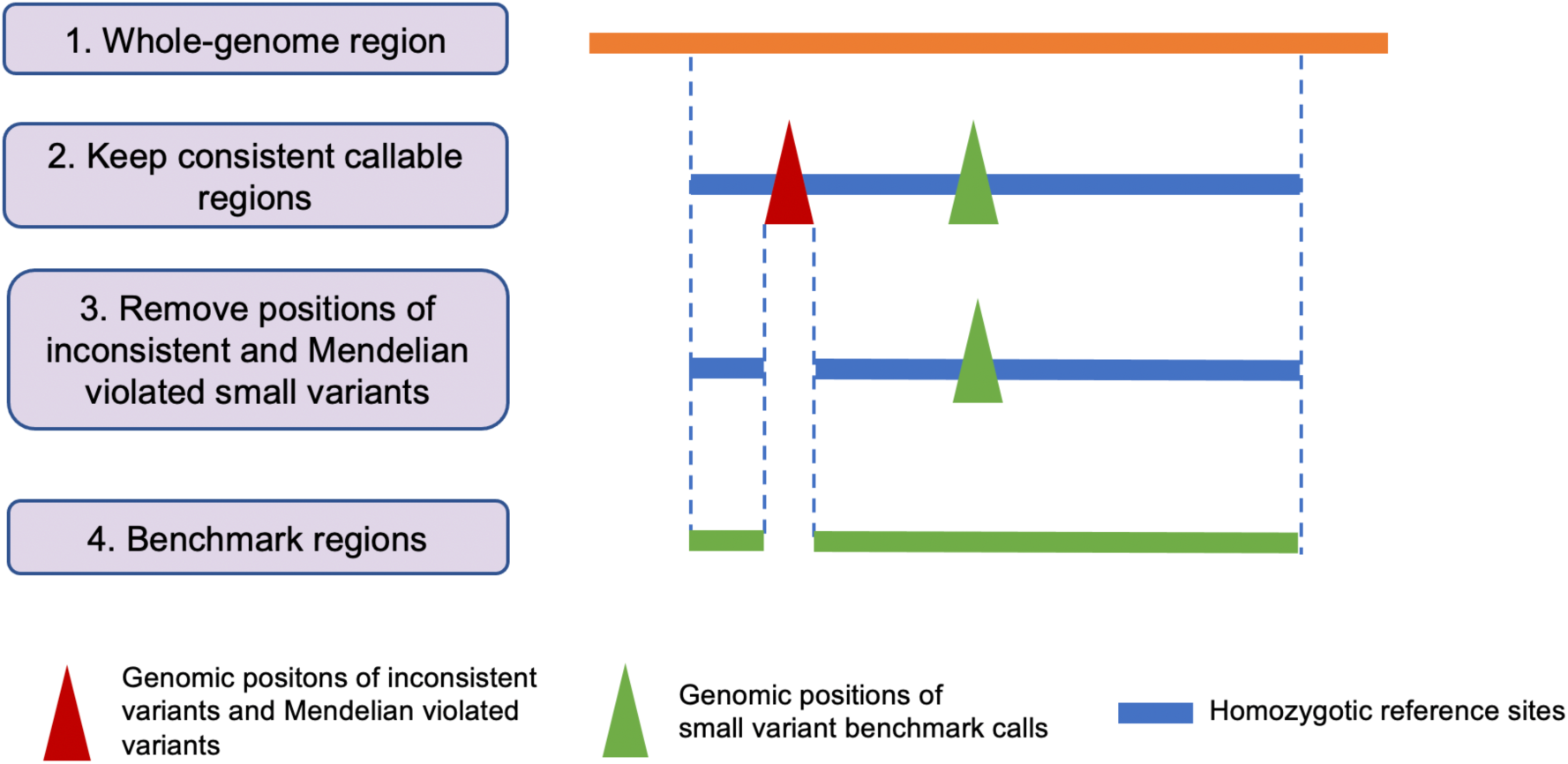
Workflow of defining benchmark regions for small variants.

**Supplementary Figure 4.**
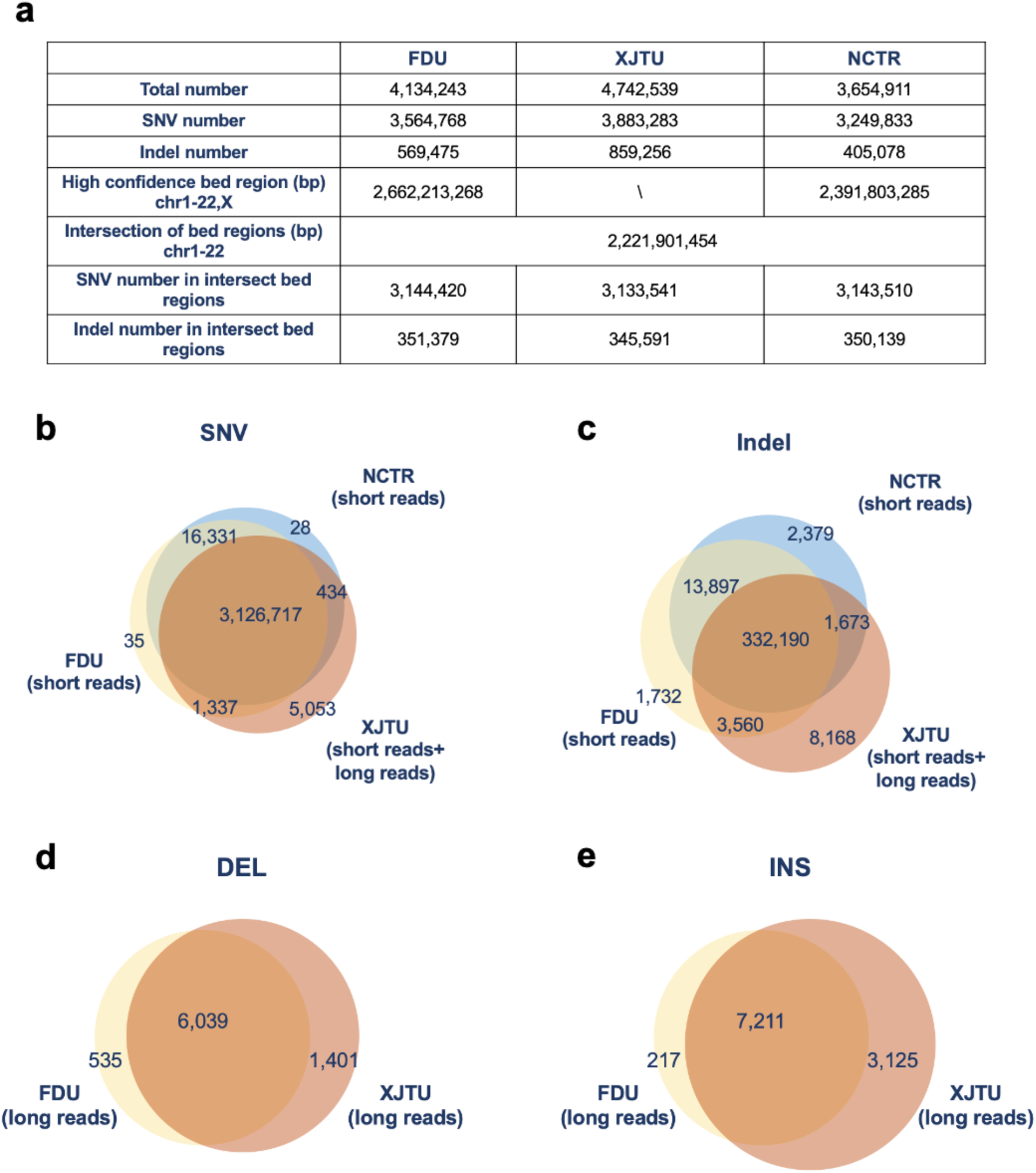
Comparison of three benchmark sets from FDU (our study), NCTR (Pan et al.), and XTJU (Jia et al.). (**a**) Statistics of small variant benchmark sets, (**b**) Venn diagram of SNV benchmark sets, (**c**) Venn diagram of indel benchmark sets, (**d**) Venn diagram of DEL benchmark sets, and (**e**) Venn diagram of INS benchmark sets.

**Supplementary Figure 5.**
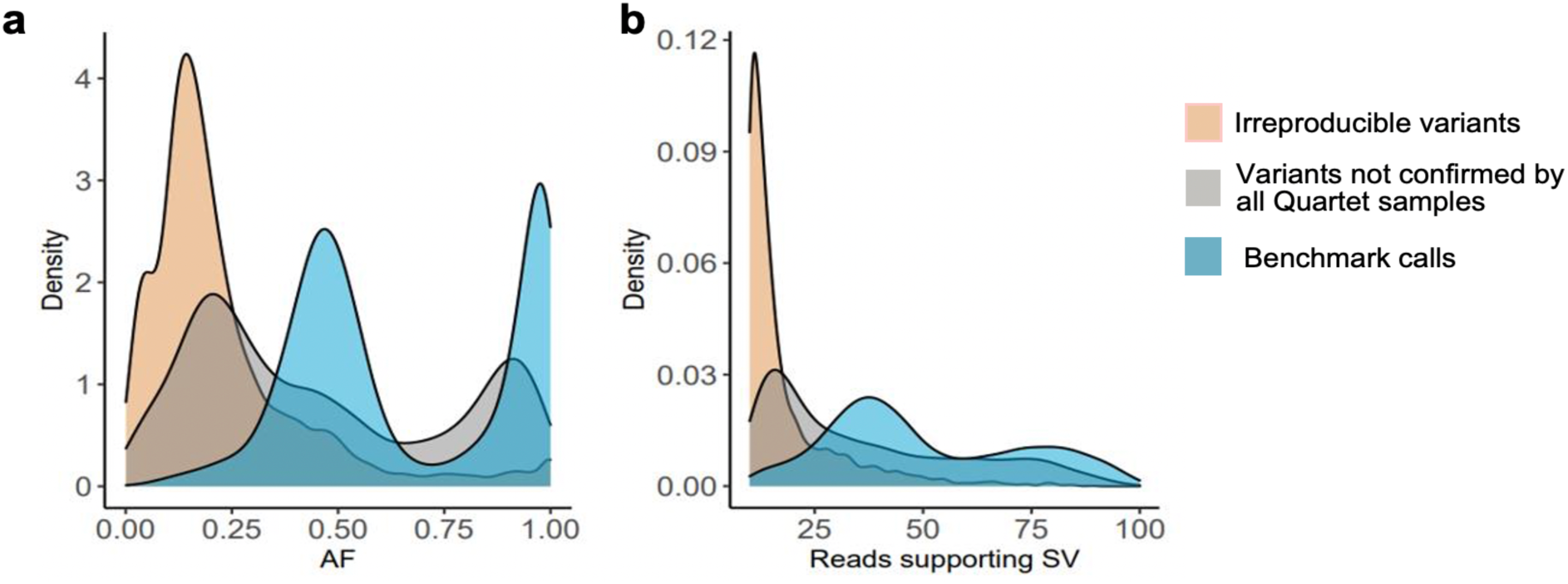
Density plots show differences in (**a**) allele frequency and (**b**) reads supporting SV, (**c** and **d**) between irreproducible SVs, reproducible but Mendelian discordant SVs and reproducible and Mendelian concordant SVs.

**Supplementary Figure 6.**
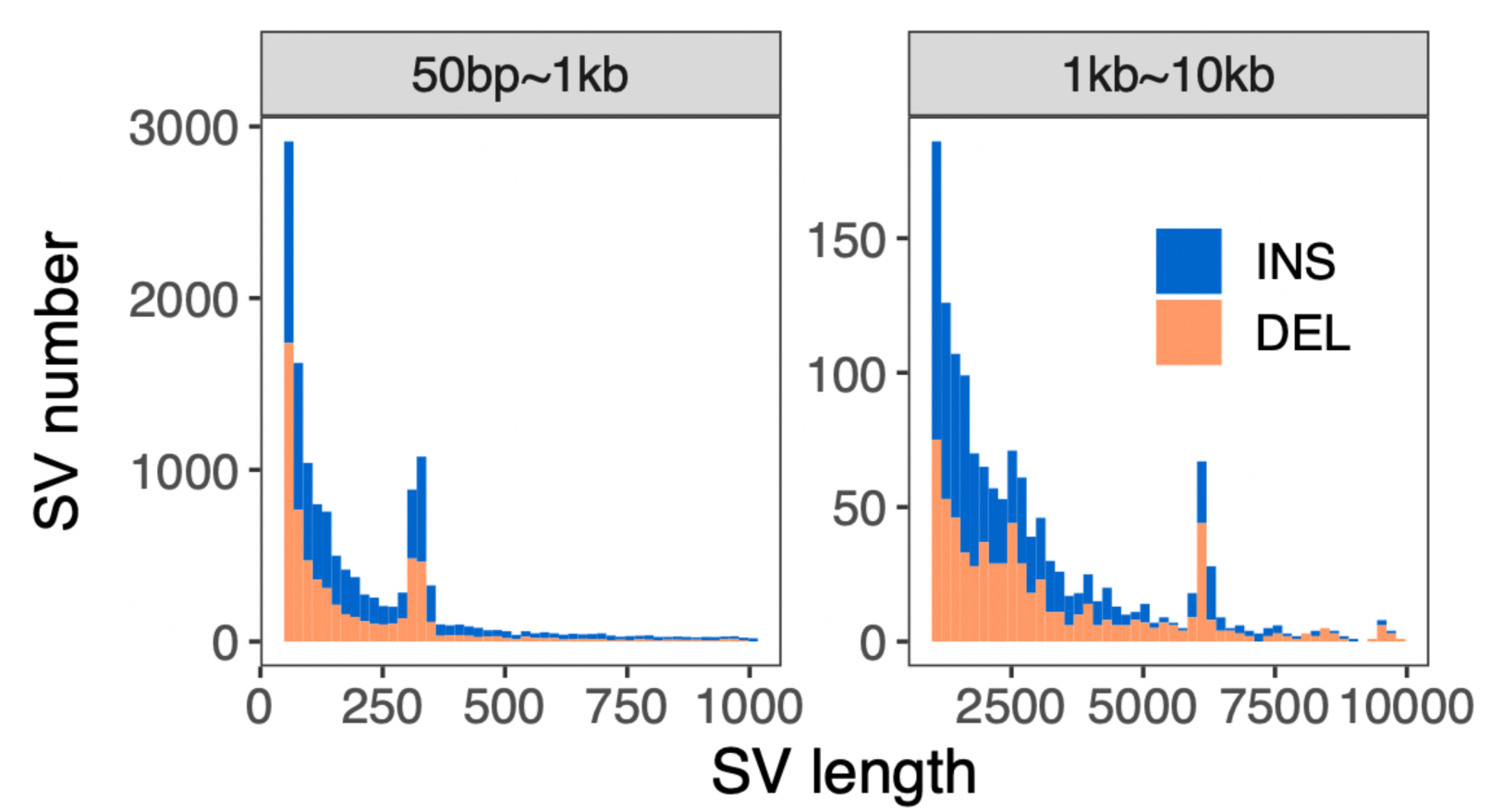
Length distribution of SV benchmark calls for the Quartet DNA reference samples.

**Supplementary Figure 7.**
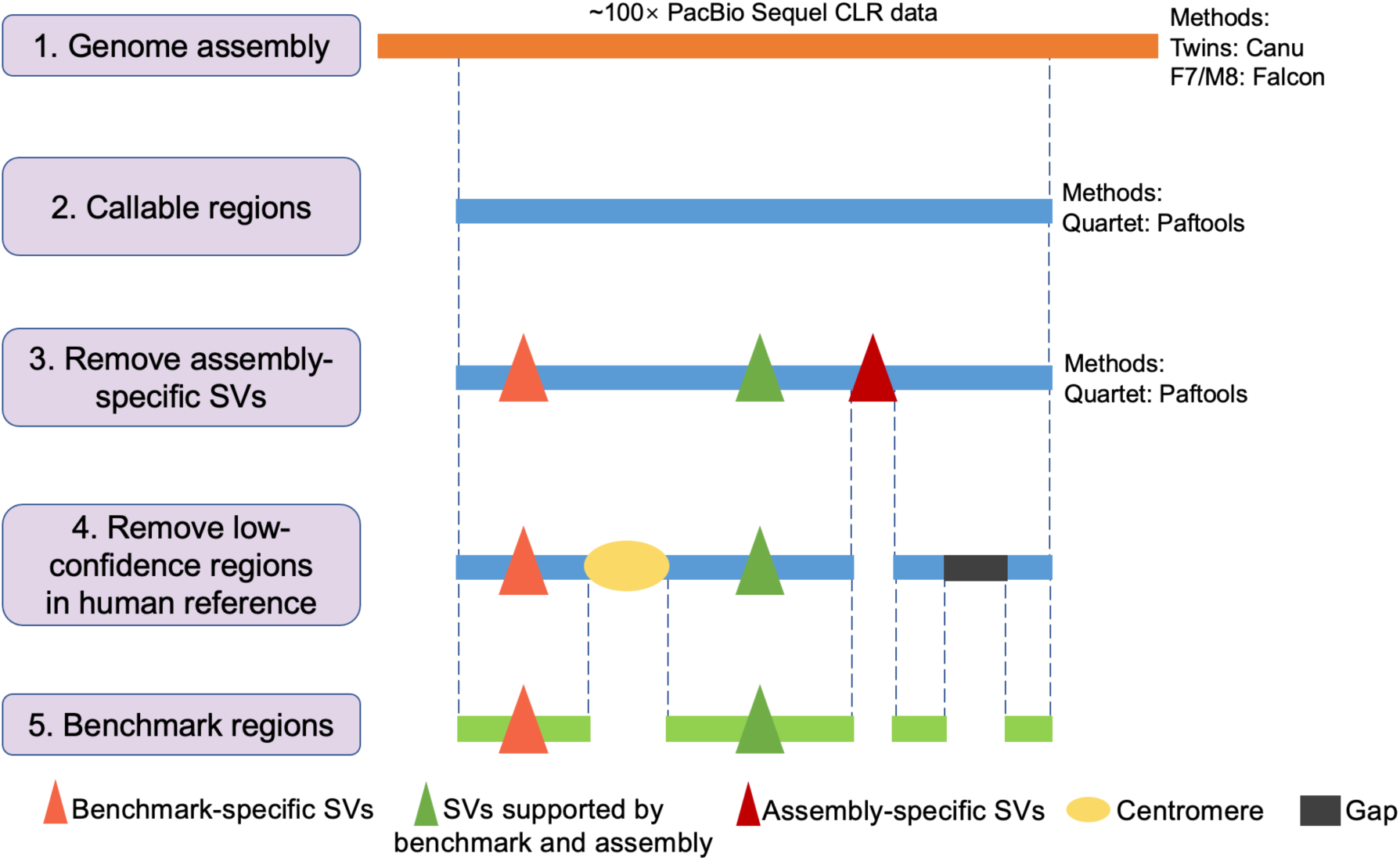
Workflow of defining benchmark regions for structural variants.

**Supplementary Figure 8.**
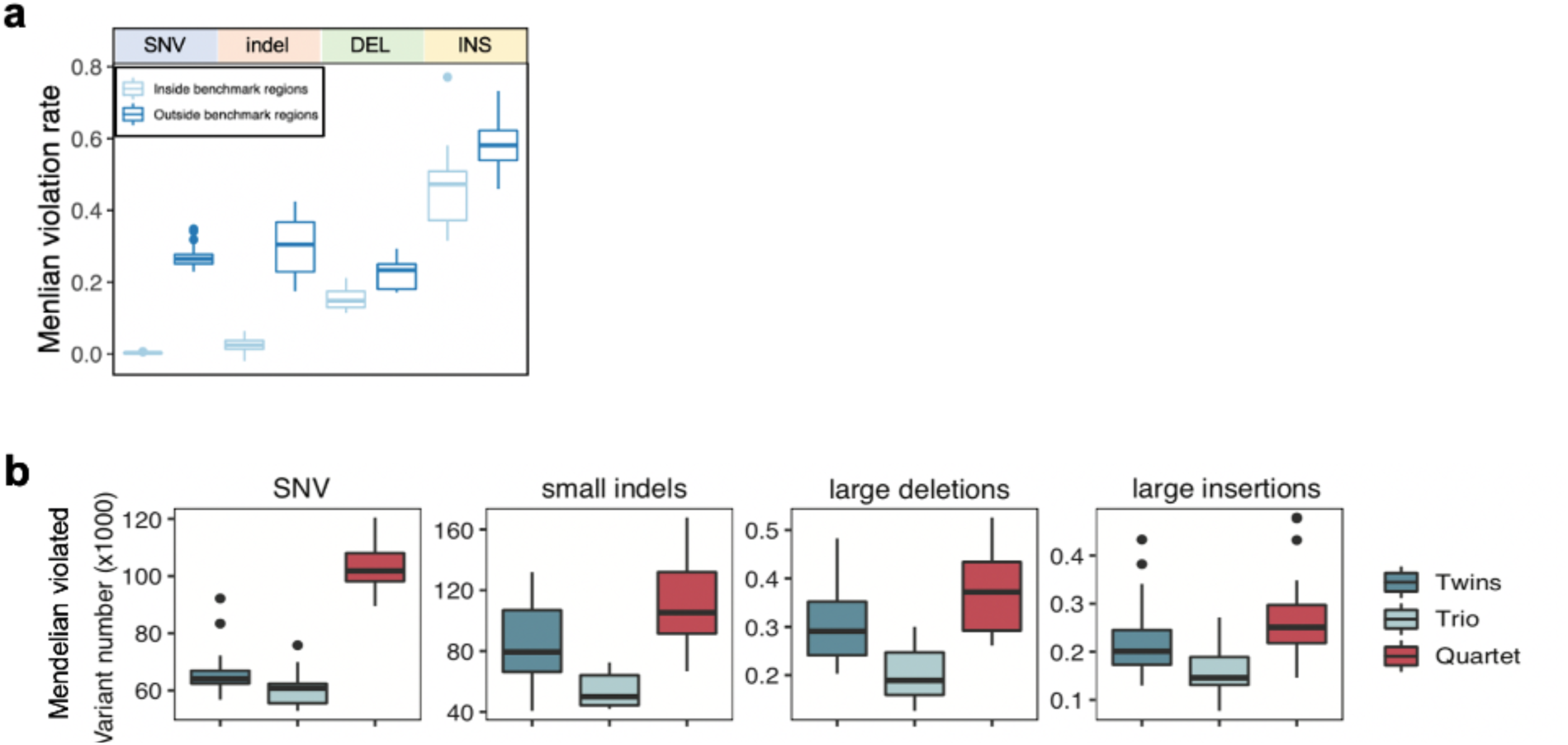
(a) Comparison of Mendelian violation rate inside and outside the benchmark regions across different variant types. **(b)** Discordant variants detected by twins (D5 and D6), Mendelian discordant variants detected by trios (D5-F7-M8 and D6-F7-M8), and Mendelian discordant variants detected by Quartet family (D5-D6-F7-M8).

**Supplementary Figure 9.**
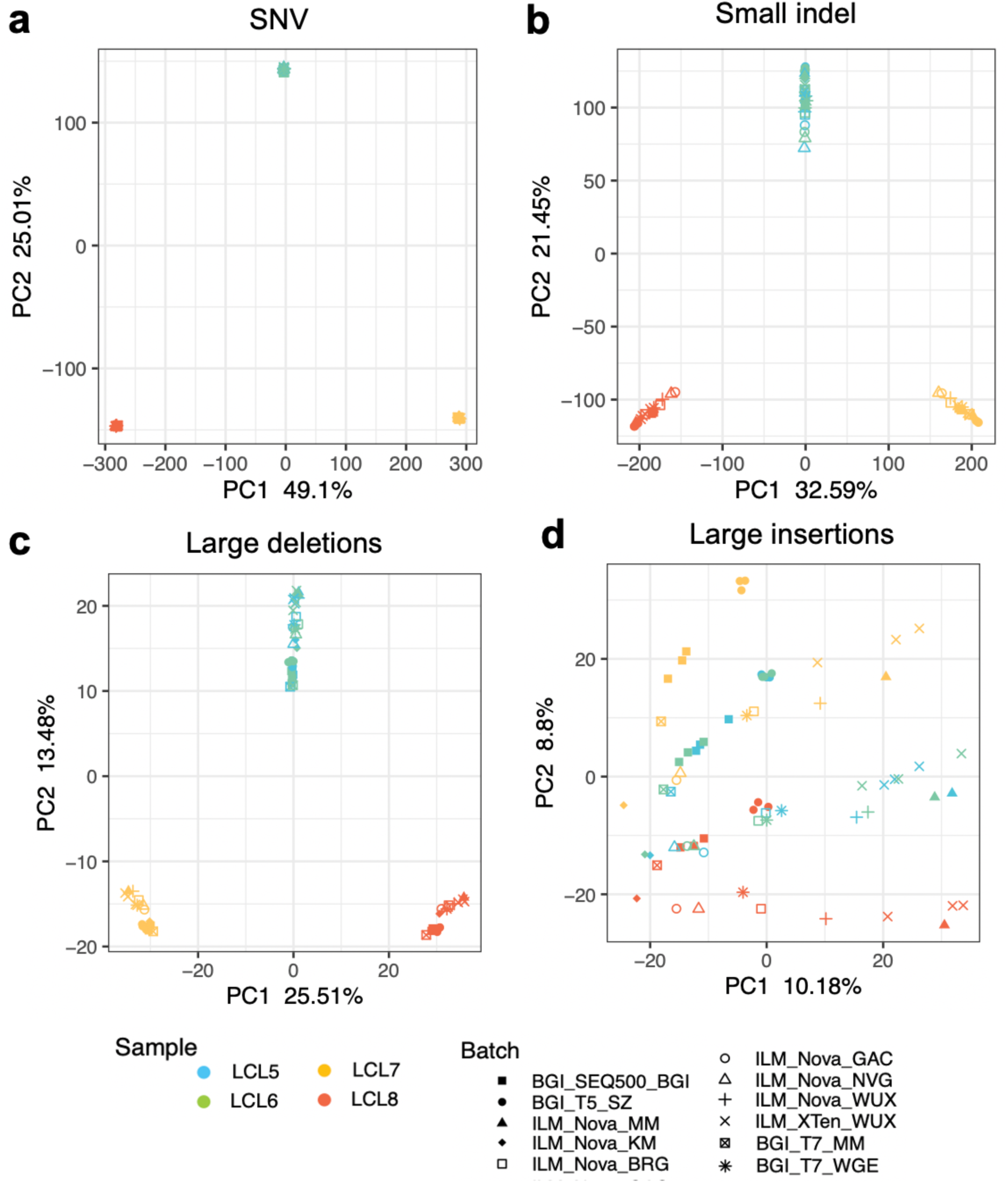
The scatterplot of the first two eigenvectors generated from PCA displayed clustering of the Quartet samples. Four different variant types from 11 batches short-read sequencing datasets are shown as PCA plots. (**a**) SNVs; (**b**) Small indels; (**c**) Large deletions; and (**d**) Large insertions.

**Supplementary Figure 10.**
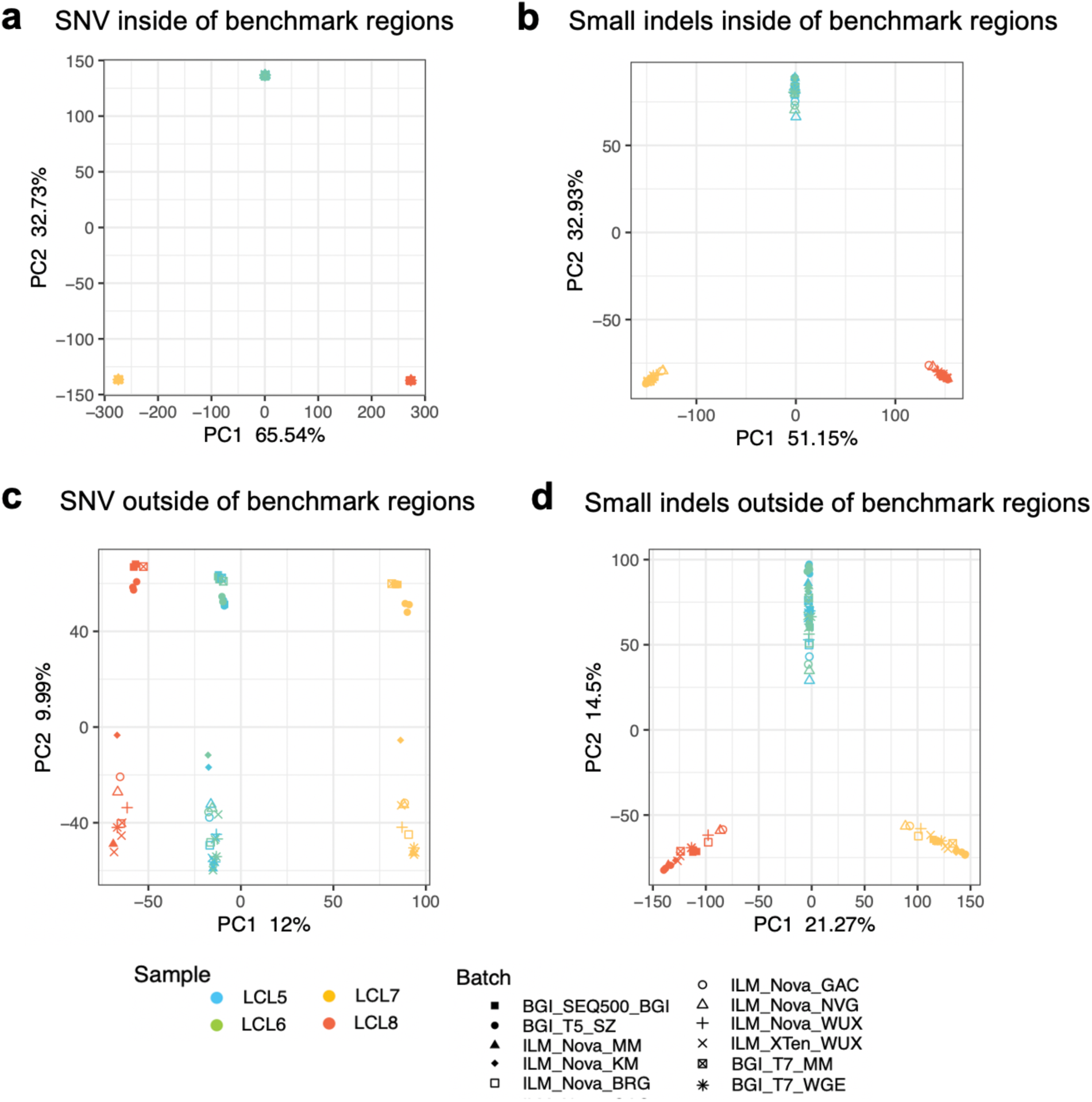
Variants called outside the benchmark regions showed more severe batch effects than variants called inside the benchmark regions. Small variants called inside and outside of benchmark regions from 11 batches of short-read sequencing datasets are shown as PCA plots. (**a**) SNVs called inside the benchmark regions; (**b**) Small indels called inside the benchmark regions; (**c**) SNVs called outside the benchmark regions; and (**d**) Small indels called outside the benchmark regions.

## Supplementary Tables

**Table 2.** Coverage of Quartet benchmark regions on coding region and clinically related genes.

| Class | Total bases (bp) | Small variant benchmark regions of Quartet<sup>1</sup> v1.1 (%) | Small variant benchmark regions of HG001 V4.2.1 (%) | SV benchmark regions of Quartet D5&D6 v1.1 (%) | SV benchmark regions of Quartet-F7 v1.1 (%) | SV benchmark regions of Quartet-M8 v1.1 (%) | SV benchmark regions of HG002 v0.6 (Tier1) (%) |
| --- | --- | --- | --- | --- | --- | --- | --- |
| GRCh38 (chr1-22, X) | 3,031,042,417 | 87.8 | 82.9 | 86.5 | 85.5 | 85.7 | 87.8 |
| Genes | 1,827,967,802 | 90.1 | 86.4 | 95.9 | 95.9 | 95.9 | 93.9 |
| Exons | 162,225,069 | 86.8 | 83.5 | 95.8 | 95.7 | 95.7 | 93.3 |
| ClinVar<sup>2</sup> | 5,713,149 | 94.4 | 88.1 | 97.6 | 97.9 | 97.0 | 94.0 |

1 The benchmark regions of the four Quartet reference samples are the same;
2 Coverage of ClinVar on the genome was used in this study.

**Supplementary Table 1.** Data from multiple short-read and long-read sequencing platforms were obtained to detect and validate small variant and structural variant benchmark calls in the Quartet reference samples.

|  | Variant type | Platforms | Library prep method | Sequencing site | Replicates | Number of reads (million) |
| --- | --- | --- | --- | --- | --- | --- |
| Discovery | Small variants | Illumina HiSeq XTen | PCR | WUX | 4 samples x 3 reps | 388.5 |
|  |  | Illumina HiSeq XTen | PCR | NVG | 4 samples x 3 reps | 328.7 |
|  |  | Illumina HiSeq XTen<sup>1</sup> | PCR | ARD | 4 samples x 3 reps | 307.9 |
|  |  | MGI-SEQ2000 | PCR | BGI | 4 samples x 3 reps | 725.3 |
|  |  | Illumina NovaSeq | PCR-free | BRG | 4 samples x 3 reps | 613.6 |
|  |  | Illumina NovaSeq | PCR-free | ARD | 4 samples x 3 reps | 581.6 |
|  |  | Illumina NovaSeq | PCR-free | ARD | 4 samples x 3 reps | 683.8 |
|  |  | Illumina NovaSeq | PCR | WUX | 4 samples x 3 reps | 327.3 |
|  |  | DNB-SEQT7 | PCR-free | WGE | 4 samples x 3 reps | 533.1 |
|  | Structural variants | Nanopore | PCR-free | GRM | 4 samples x 1 rep | 26.8 |
|  |  | PacBio Sequel CLR | PCR-free | NOM | 4 samples x 1 rep | 33 |
|  |  | PacBio Sequel II CLR | PCR-free | ARD | 4 samples x 1 rep | 5.3 |
| Validation | Small variants | PacBio Sequel II CCS<sup>2</sup> | PCR-free | GRM | 4 samples x 1 rep | 11.7 |
|  |  | PMRA | - | BIO | 4 samples x 16 reps | - |
|  | Structural variants | 10x Genomics | PCR | BIO | 4 samples x 1 rep | 320 |
|  |  | BioNano | PCR-free | NOM | 4 samples x 1 rep | - |
|  |  | PacBio Sequel II CCS<sup>2</sup> | PCR-free | GRM | 4 samples x 1 rep | 11.7 |
|  |  | Illumina HiSeq XTen<sup>1</sup> | PCR | ARD | 4 samples x 3 reps | 307.9 |
| <sup>1</sup> The same short-read WGS datasets used to construct small variant benchmark calls and validate structural variants benchmark calls; |  |  |  |  |  |  |
| <sup>2</sup> The same PacBio CCS datasets used to validate both small variant and structural variant benchmark calls. |  |  |  |  |  |  |

**Supplementary Table 2.** Mapping and calling statistics of short-read sequencing datasets. See attachment

**Supplementary Table 3.** Statistics of long-read raw sequencing datasets. See attachment

**Supplementary Table 4.** Mapping statistics of long-read sequencing datasets. See attachment

**Supplementary Table 5.** Statistics of *de novo* and somatic small variants for the Quartet. See attachment

**Supplementary Table 6.**
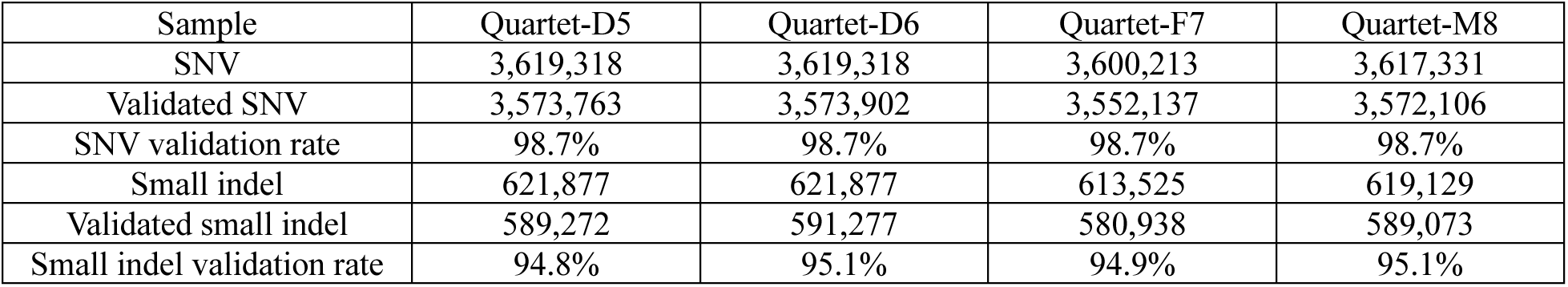
Validation of small variants by PacBio CCS, which are reproducible among call sets and Mendelian consistent in the Quartet family.

**Supplementary Table 7.** Validation of small variants by PMRA, which are reproducible among call sets and Mendelian consistent in the Quartet family.

| Sample | Quartet-D5/D6 | Quartet-F7 | Quartet-M8 |
| --- | --- | --- | --- |
| PMRA consistent sites | 793,024 | 793,024 | 793,024 |
| Validated sites | 790,293 | 790,274 | 790,290 |
| Validation rate | 99.7% | 99.7% | 99.7% |

**Supplementary Table 8.** Statistics of *de novo* SVs for the Quartet family.

| Class | Chr | Start | End | SV type | Size | F7 | M8 | D5 | D6 |
| --- | --- | --- | --- | --- | --- | --- | --- | --- | --- |
| Twin-shared | chr14 | 105,982,942 | 105,984,761 | DEL | 1,819 | 0/0 | 0/0 | 0/1 | 0/1 |
|  | chr14 | 105,989,116 | 105,989,116 | INS | 67 | 0/0 | 0/0 | 0/1 | 0/1 |
| Twin-specific | chr12 | 85,821,376 | 85,821,376 | INS | 55 | 0/0 | 0/0 | 0/0 | 0/1 |
|  | chr7 | 2,492,258 | 2,492,347 | DEL | 89 | 0/0 | 0/0 | 0/1 | 0/0 |
|  | chr8 | 86,407,355 | 86,409,907 | DEL | 2,552 | 0/0 | 0/0 | 0/1 | 0/0 |
|  | chrX | 129,048,461 | 129,048,537 | DEL | 76 | 0/0 | 0/0 | 0/0 | 0/1 |

**Supplementary Table 9.**
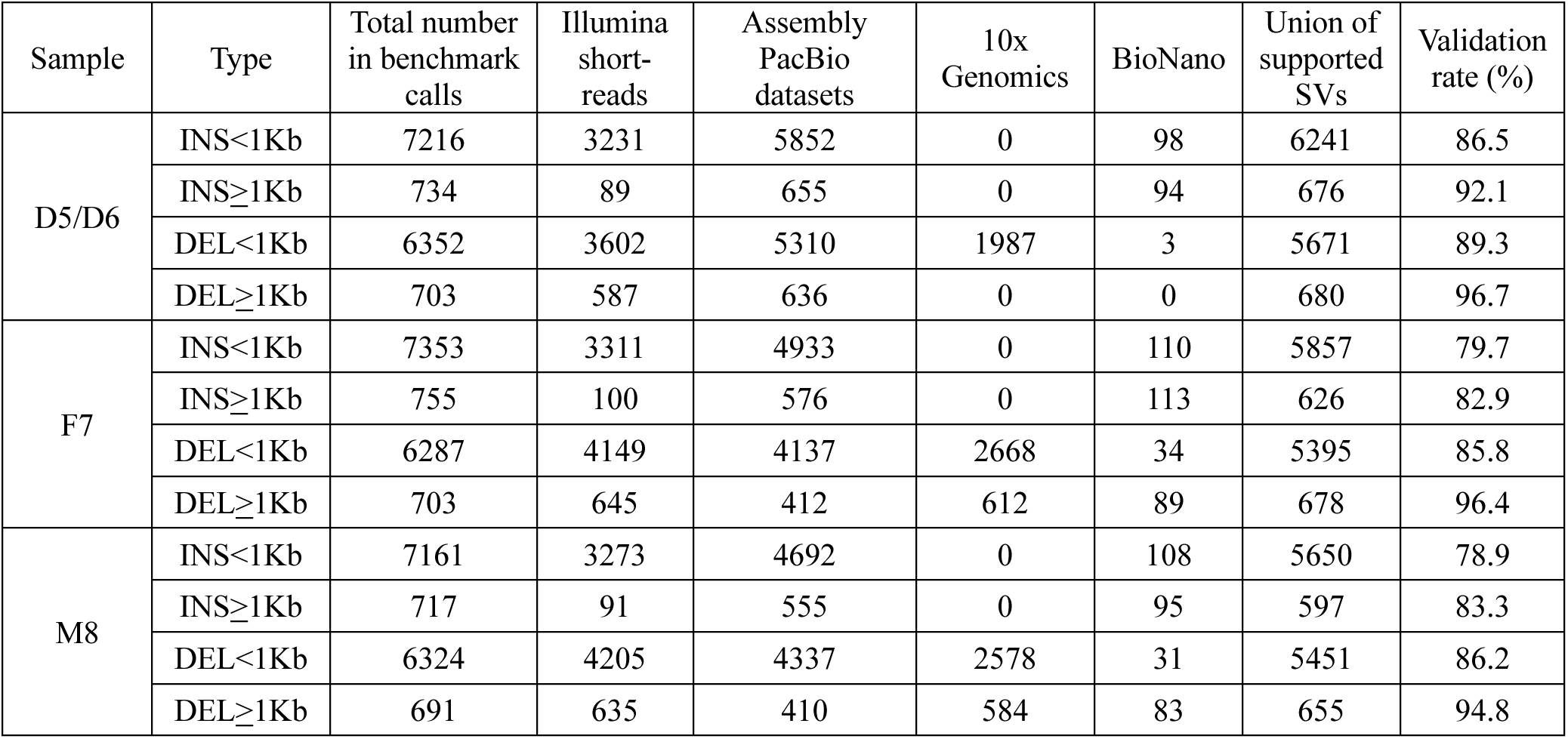
Validation of SV benchmark calls by Illumina short-reads, 10x Genomics, BioNano, and assembly PacBio reads.

**Supplementary Table 10.** Datasets for proficiency test analysis and batch effect analysis.

| Batch | Library preparation method | Sequencing platforms | Sequencing labs | Replicates of the four Quartet samples | Median coverage | Median insert size |
| --- | --- | --- | --- | --- | --- | --- |
| 1 | PCR | Illumina NovaSeq 6000 | BRG | 1 | 34x | 321 |
| 2 | PCR | Illumina NovaSeq 6000 | GAC | 1 | 27x | 279 |
| 3 | PCR | Illumina NovaSeq 6000 | NVG | 1 | 25x | 290 |
| 4 | PCR | Illumina NovaSeq 6000 | WUX | 1 | 26x | 412 |
| 5 | PCR | BGI-SEQ500 | BGI | 3 | 40x | 234 |
| 6 | PCR | Illumina XTen | WUX | 3 | 35x | 396 |
| 7 | PCR-free | DNB-SEQT5 | BGI | 3 | 50x | 350 |
| 8 | PCR-free | Illumina NovaSeq 6000 | KM | 1 | 33x | 206 |
| 9 | PCR-free | Illumina NovaSeq 6000 | WUX | 1 | 30x | 401 |
| 10 | PCR-free | DNB-SEQT7 | WUX | 1 | 30x | 259 |
| 11 | PCR-free | DNB-SEQT7 | WGE | 1 | 35x | 359 |

